# Thymocytes trigger self-antigen-controlling pathways in immature medullary thymic epithelial stages

**DOI:** 10.1101/2020.11.29.399923

**Authors:** Noëlla Lopes, Nicolas Boucherit, Jonathan Charaix, Pierre Ferrier, Matthieu Giraud, Magali Irla

## Abstract

Interactions of developing T cells with Aire^+^ medullary thymic epithelial cells expressing high levels of MHCII molecules (mTEC^hi^) are critical for the induction of central tolerance. In turn, thymocytes regulate the cellularity of Aire^+^ mTEC^hi^. However, it remains unknown whether thymocytes control Aire^+^ mTEC^hi^-precursors that are contained in mTEC^lo^ cells or other mTEC^lo^ subsets that have recently been delineated or identified by single-cell transcriptomic analyses. Here, using three distinct transgenic mouse models, in which antigen-presentation between mTECs and CD4^+^ thymocytes is perturbed, we show by high-throughput RNA-seq that self-reactive CD4^+^ thymocytes induce in mTEC^lo^ the expression of tissue-restricted self-antigens, cytokines, chemokines and adhesion molecules important for T-cell development. This gene activation program is combined with a global increase of the active H3K4me3 histone mark. Finally, we show that these interactions induce key mTEC transcriptional regulators and govern mTEC^lo^ subset composition, including Aire^+^ mTEC^hi^-precursors, post-Aire and tuft-like mTECs. Our genome-wide study thus reveals that self-reactive CD4^+^ thymocytes control multiple unsuspected facets from immature stages of mTECs, which determines their heterogeneity.

## Introduction

The thymic medulla ensures the generation of a self-tolerant T-cell repertoire (Klein et al, 2014; Lopes et al, 2015). By their unique ability to express tissue-restricted self-antigens (TRAs) (Derbinski et al, 2001; Sansom et al, 2014), medullary thymic epithelial cells (mTECs) promote the development of Foxp3^+^ regulatory T cells and the deletion by apoptosis of self-reactive thymocytes capable to induce autoimmunity (Klein et al, 2019). The expression of TRAs that mirrors body’s self-antigens is controlled by Aire (*Autoimmune regulator*) and Fezf2 (*Fez family zinc finger 2*) transcription factors (Anderson et al, 2002; Takaba et al, 2015). Aire-dependent TRAs are generally characterized by a repressive chromatin state enriched in the trimethylation of lysine-27 of histone H3 (H3K27me3) histone mark (Handel et al, 2018; Org et al, 2009; Sansom et al, 2014). In accordance with their essential role in regulating the expression of TRAs, *Aire*^-/-^ and *Fezf2*^-/-^ mice show defective clonal deletion of autoreactive thymocytes and develop signs of autoimmunity in several peripheral tissues (Anderson et al, 2002; Takaba et al, 2015).

Based on the level of the co-expressed MHC class II and CD80 molecules, mTECs were initially subdivided into mTEC^lo^ (MHCII^lo^CD80^lo^) and mTEC^hi^ (MHCII^hi^CD80^hi^) (Gray et al, 2006). The relationship between these two subsets has been established with reaggregate thymus organ cultures in which mTEC^lo^ give rise to mature Aire^+^ mTEC^hi^ (Gabler et al, 2007; Gray et al, 2007). Although mTEC^hi^ express a highly diverse array of TRAs under Aire’s action that releases stalled RNA polymerase and modulates chromatin accessibility, mTEC^lo^ already express a substantial amount of TRAs (Derbinski et al, 2005; Giraud et al, 2012; Koh et al, 2018; Kyewski & Klein, 2006; Sansom et al, 2014). Recent single-cell advances indicate that the heterogeneity of mTECs, especially in the mTEC^lo^ compartment, is more complex than previously thought (Kadouri et al, 2019). mTEC^lo^ with intermediate levels of CD80 (CD80^int^) lie into mTEC clusters that were defined as proliferating and maturational, expressing Fezf2 and preceding Aire^+^ mTEC^hi^ (Baran-Gale et al, 2020; Dhalla et al, 2020). On the other hand, mTEC^lo^ with low or no expression of CD80 (CD80^neg/lo^) have been shown to be divided into three main subsets: CCL21^+^ mTECs (Lkhagvasuren et al, 2013), involucrin^+^TPA^hi^ post-Aire mTECs corresponding to the ultimate mTEC differentiation stage (Metzger et al, 2013; Michel et al, 2017; Nishikawa et al, 2010) and the newly reported tuft-like mTECs that show properties of gut chemosensory epithelial tuft cells expressing the doublecortin-like kinase 1 (DCLK1) marker (Bornstein et al, 2018; Miller et al, 2018).

In the postnatal thymus, while mTECs control the selection of thymocytes, conversely CD4^+^ thymocytes control the cellularity of Aire^+^ mTEC^hi^ by activating RANK and CD40 signaling pathways (Akiyama et al, 2008; Hikosaka et al, 2008; Irla et al, 2008). These bidirectional interactions between mTECs and thymocytes are commonly referred to as thymic crosstalk (Lopes et al, 2015; van Ewijk et al, 1994). However, it remains unknown whether CD4^+^ thymocytes act exclusively on Aire^+^ mature mTEC^hi^ or upstream on their precursors contained in mTEC^lo^ and whether the development of the newly identified Fezf2^+^, post-Aire and tuft-like subsets is regulated or not by CD4^+^ thymocytes.

In this study, using high-throughput RNA-sequencing (RNA-seq), we show that self-reactive CD4^+^ thymocytes upregulate in mTEC^lo^ the expression of TRAs, chemokines, cytokines and adhesion molecules involved in T-cell development. This gene activation program correlates with increased levels of the active trimethylation of lysine-4 of histone 3 (H3K4me3) mark, particularly in the loci of Fezf2-dependent and Aire/Fezf2-independent TRAs, indicative of an epigenetic regulation for their expression. Our data also reveal that self-reactive CD4^+^ thymocytes induce in mTEC^lo^ key transcriptional regulators pivotal for mTEC differentiation and function. Accordingly, self-reactive CD4^+^ thymocytes control the composition of the mTEC^lo^ compartment, *i.e.*: Aire^+^ mTEC^hi^-precursors, post-Aire cells and tuft-like mTECs. Finally, we demonstrate that disrupted MHCII/TCR interactions between mTECs and CD4^+^ thymocytes leads to an altered TCRVβ repertoire in mature T cells containing self-specificities capable of inducing multi-organ autoimmunity. Altogether, our genome-wide study reveals that self-reactive CD4^+^ thymocytes control the developmental transcriptional programs in mTEC^lo^, which condition their differentiation and function as inducers of T-cell tolerance.

## Results

### CD4^+^ thymocytes induce key transcriptional programs in mTEC^lo^ cells

Several NF-κb members are involved in Aire^+^ mTEC^hi^ development (Burkly et al, 1995; Lomada et al, 2007; Riemann et al, 2017; Shen et al, 2019; Zhang et al, 2006). However, it remains unclear whether the NF-κb pathways or other signaling pathways are activated by CD4^+^ thymocytes specifically in mTEC^lo^ cells. To investigate the effects of CD4^+^ thymocytes in mTEC^lo^, we used mice deficient in CD4^+^ thymocytes (ΔCD4 mice) because they lack the promoter IV of the class II transactivator (*Ciita*) gene that controls MHCII expression in cortical TECs (cTECs) (Waldburger et al, 2003). We first analyzed by flow cytometry the total and phosphorylated forms of IKKα, p65 and RelB NF-κb members and p38 and Erk1/2 MAPK proteins in mTEC^lo^ from ΔCD4 mice according to the gating strategy shown in **Figure S1A**. Interestingly, the phosphorylation level of IKKα and p38 MAPK was substantially reduced in ΔCD4 mice **(Figure 1A,B; Figure S2)**, indicating that CD4^+^ thymocytes activate in mTEC^lo^ the IKKα intermediate of non-classical NF-κB pathway and the p38 MAPK pathway.

**Figure 1.**
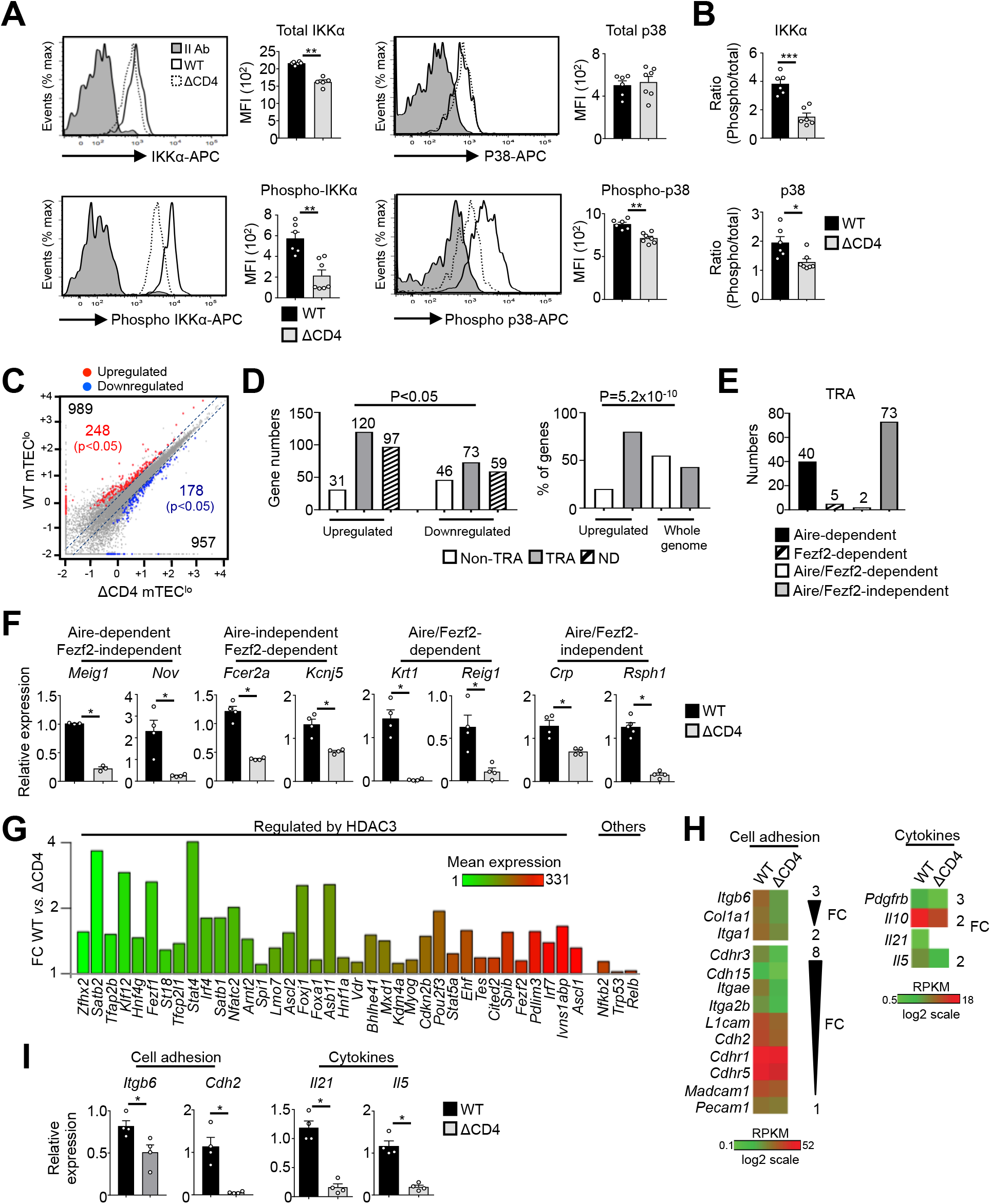
The transcriptional profile and IKKα and p38 MAPK signaling pathways are impaired in mTEC^lo^ of ΔCD4 mice. **(A,B)** Total IKKα, p38 MAPK, phospho-IKKα(Ser180)/IKKβ(Ser181) and p38 MAPK (Thr180/Tyr182) **(A)** and the ratio of phospho/total proteins **(B)** analyzed by flow cytometry in mTEC^lo^ from WT and ΔCD4 mice. Data are representative of 2 independent experiments (n=3-4 mice per group and experiment). **(C)** Scatter-plot of gene expression levels (FPKM) of mTEC^lo^ from WT *versus* ΔCD4 mice. Genes with fold difference ≥2 and p-adj <0.05 were considered as upregulated or downregulated genes (red and blue dots, respectively). RNA-seq was performed on 2 independent biological replicates with mTEC^lo^ derived from 3-5 mice. **(D)** Numbers of TRAs and non-TRAs in genes up- and downregulated (left panel) and the proportion of upregulated TRAs compared to those in the all genome (right panel). ND: not determined. **(E)** Numbers of induced Aire-dependent, Fezf2-dependent, Aire/Fezf2-dependent and Aire/Fezf2-independent TRAs. **(F)** The expression of Aire-dependent (*Meig1, Nov*), Fezf2-dependent (*Fcer2a, Kcnj5*), Aire/Fezf2-dependent (*Krt1, Reig1*) and Aire/Fezf2-independent (*Crp, Rsph1*) TRAs measured by qPCR in WT (n=3-4) and ΔCD4 (n=3-4) mTEC^lo^. **(G)** Expression fold change in HDAC3-induced transcriptional regulators and other transcription factors in WT *versus* ΔCD4 mTEC^lo^. The color code represents gene expression level. **(H)** Heatmaps of selected genes encoding for cell adhesion molecules and cytokines. **(I)** *Itgb6, Cdh2, Il21* and *Il5* mRNAs were measured by qPCR in WT (n=4) and ΔCD4 (n=4) mTEC^lo^. Error bars show mean◻±◻SEM, *p<◻0.05, **p<0.01, ***p<0.001 using two-tailed Mann-Whitney test for **A,B,F,I** and Chi-squared test for **D**.

To gain insights into the effects of CD4^+^ thymocytes in mTEC^lo^, we analyzed by high-throughput RNA-seq the gene expression profiles of mTEC^lo^ purified from WT and ΔCD4 mice **(Figure S1B).** We found that CD4^+^ thymocytes upregulated 989 genes (fold-change FC>2) reaching significance for 248 of them (Cuffdiff p<0.05) **(Figure 1C)**. 957 genes were also downregulated (FC<0.5) with 178 genes reaching significance (Cuffdiff p<0.05). We analyzed whether the genes significantly up- or downregulated by CD4^+^ thymocytes corresponded to TRAs, as defined by an expression restricted to one to five of peripheral tissues (Sansom et al, 2014). Interestingly, the genes upregulated by CD4^+^ thymocytes exhibited ~4-fold more of TRAs over non-TRAs **(Figure 1D, left panel)**. The comparison of the proportion of TRAs among the upregulated genes with those of the genome revealed a strong statistical TRA overrepresentation (p=5.2×10^-10^) **(Figure 1D, right panel)**. Most of the TRAs upregulated by CD4^+^ thymocytes are sensitive to the action of Aire (Aire-dependent TRAs) or controlled by Aire and Fezf2-independent mechanisms (Aire/Fezf2-independent TRAs) **(Figure 1E and Table S1).** The upregulation of some of these TRAs by CD4^+^ thymocytes was confirmed by qPCR in mTEC^lo^ purified from ΔCD4 and MHCII^-/-^ mice, the latter also lacking CD4^+^ thymocytes **(Figure 1F and Figure S3A)**.

Remarkably, among the non-TRAs upregulated by CD4^+^ thymocytes in mTEC^lo^, 37 corresponded to 50 mTEC-specific transcription factors that are induced by the histone deacetylase 3 (HDAC3) (Goldfarb et al, 2016) **(Figure 1G)**. Some of them, such as the interferon regulatory factor 4 (*Irf4*), *Irf7* and the Ets transcription factor member, *Spib*, have been described to regulate mTEC differentiation and function (Akiyama et al, 2014; Haljasorg et al, 2017; Otero et al, 2013). We also identified other transcription factors such as *Nfkb2*, *Trp53* and *Relb* implicated in mTEC differentiation (Riemann et al, 2017; Rodrigues et al, 2017; Zhang et al, 2006). Finally, we found that CD4^+^ thymocytes control in mTEC^lo^ the expression of some cytokines and cell adhesion molecules such as integrins and cadherins **(Figure 1H,I)**. Altogether, these results provide the first evidence that CD4^+^ thymocytes are able to induce in mTEC^lo^ essential transcriptional regulators for mTEC differentiation and function but also the expression of TRAs, adhesion molecules and cytokines implicated in thymocyte selection.

### CD4^+^ thymocytes regulate maturational programs in mTEC^lo^ through MHCII/TCR interactions

We next investigated by which mechanism CD4^+^ thymocytes regulate the transcriptional programs in mTEC^lo^. Given that MHCII/TCR interactions with mTECs are critical for CD4^+^ T-cell selection (Klein et al, 2019), we hypothesized that they could play an important role in initiating transcriptional programs that govern the functional and developmental properties of mTEC^lo^. To this end, we used a unique transgenic mouse model in which MHCII expression is selectively abrogated in mTECs (mTEC^ΔMHCII^ mice) (Irla et al, 2008). In contrast to their WT counterparts, we found that OVA_323-339_-loaded mTECs from mTEC^ΔMHCII^ mice were ineffective at activating OTII-specific CD4^+^ T cells, demonstrating that antigen-presentation ability of mTECs to CD4^+^ T cells is impaired in these mice **(Figure 2A).**

**Figure 2.**
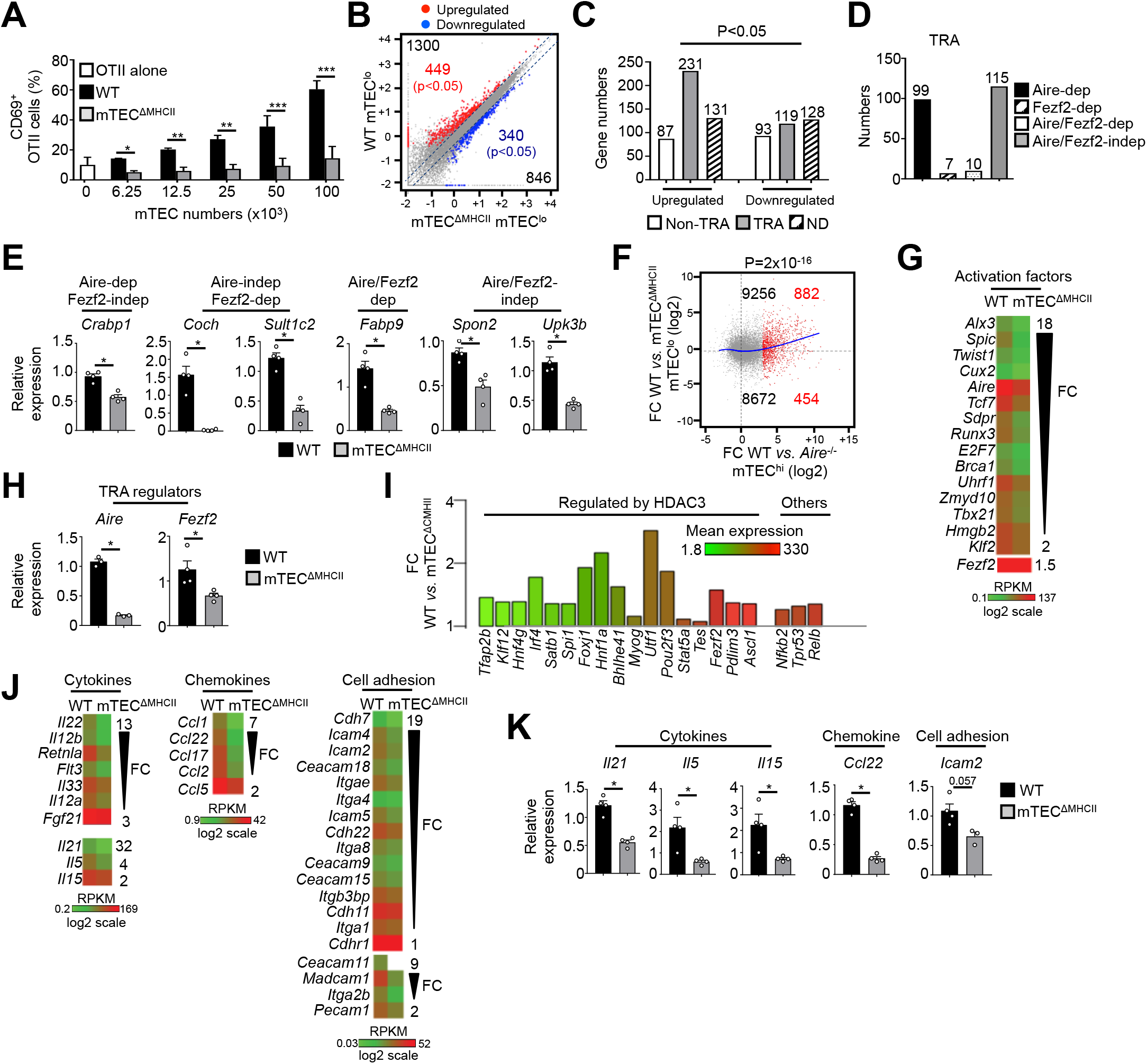
The transcriptional and functional properties of mTEC^lo^ are impaired in mTEC^ΔMHCII^ mice. **(A)** Percentages of CD69^+^ OTII CD4^+^ T cells cultured or not with variable numbers of OVA_323-339_-loaded WT or mTEC^ΔMHCII^ mTECs derived of 2 independent experiments (n=2-3 mice per group and experiment). **(B)** Scatter-plot of gene expression levels (FPKM) of mTEC^lo^ from WT *versus* mTEC^ΔMHCII^ mice. Genes with fold difference ≥2 and p-adj <0.05 were considered as upregulated or downregulated genes (red and blue dots, respectively). RNA-seq was performed on 2 independent biological replicates with mTEC^lo^ derived from 3-5 mice. **(C)** Numbers of TRAs and non-TRAs in genes up- and downregulated in mTEC^lo^ from WT *versus* mTEC^ΔMHCII^ mice. ND: not determined. **(D)** Numbers of induced TRAs regulated or not by Aire and/or Fezf2. **(E)** Aire-dependent (*Crabp1*), Fezf2-dependent (*Coch, Sult1c2*), Aire/Fezf2-dependent (*Fabp9*) and Aire/Fezf2-independent (*Spon2, Upk3b*) TRAs were measured by qPCR in mTEC^lo^ from WT (n=4) and mTEC^ΔMHCII^ (n=4) mice. **(F)** Scatter-plot of gene expression variation in mTEC^lo^ from WT *versus* mTEC^ΔMHCII^ mice and in mTEC^hi^ from WT *versus* Aire^-/-^ mice. The loess fitted curve is shown in blue and the induced Aire-dependent genes (FC>5) in red. **(G)** Heatmap of selected activation factors. **(H)** *Aire* and *Fezf2* mRNAs were measured by qPCR in mTEC^lo^ from WT (n=3-4) and mTEC^ΔMHCII^ (n=4) mice. **(I)** Fold change in the expression of HDAC3-induced transcriptional regulators and other transcription factors in WT *versus* mTEC^ΔMHCII^ mice. The color code represents gene expression level. **(J)** Heatmap of selected cytokines, chemokines and cell adhesion molecules. **(K)** *Il21*, *Il5, Il15, Ccl22* and *Icam2* mRNAs were measured by qPCR in mTEC^lo^ from WT (n=4) and mTEC^ΔMHCII^ (n=3-4) mice. Error bars show mean◻±◻SEM, *p<0.05, **p<0.01, ***p<0.001 using two-tailed Mann-Whitney test for **A,E,H,K** and Chi-squared test for **C,F**.

The comparison of the gene expression profiles of mTEC^lo^ purified from WT and mTEC^ΔMHCII^ mice **(Figure S1B)** revealed that MHCII/TCR interactions with CD4^+^ thymocytes resulted in the upregulation of 1300 genes (FC>2), 449 of them reaching statistical significance (Cuffdiff p<0.05). 846 genes were also downregulated (FC<0.5) with 340 reaching significance (Cuffdiff p<0.05) **(Figure 2B).** Similarly to the comparison of WT versus ΔCD4 mice **(Figure 1D)**, the genes significantly upregulated by MHCII/TCR interactions in mTEC^lo^ corresponded preferentially to TRAs (p=4.5×10^-13^) that are mainly Aire-dependent and Aire/Fezf2-independent **(Figures 2C-E and Table S2)**. In line with the recent discovery of Aire expression in the fraction of mTEC^lo^ that express intermediate levels of CD80 and which are found in the proliferating and maturational stage mTEC single-cell clusters (Dhalla et al, 2020), we found a strong correlation (p=2×10^-16^) between gene upregulation induced by MHCII/TCR interactions and the responsiveness of genes to Aire’s action obtained from the comparison between WT and *Aire*^-/-^ mTEC^hi^ **(Figure 2F).** These data are in agreement with the identification of a list of activation factors including *Aire* among of the non-TRA genes induced by MHCII/TCR interactions with CD4^+^ thymocytes in mTEC^lo^ **(Figure 2G).** WT mTEC^lo^ expressed ~4.5-fold more *Aire* compared to mTEC^lo^ from mTEC^ΔMHCII^ mice with substantial levels of 73.7 and 15.8 FPKM, respectively. For comparison *Aire* expression level in WT mTEC^hi^ was 448.9 FPKM. WT mTEC^lo^ also expressed ~1.5-fold more *Fezf2* compared to mTEC^lo^ from mTEC^ΔMHCII^ mice. This upregulation in *Aire* and *Fezf2* was also confirmed by qPCR **(Figure 2H).** These results highlight the importance of MHCII/TCR interactions with CD4^+^ thymocytes in upregulating *Aire* and *Fezf2* mRNAs and some of their associated TRAs in mTEC^lo^. Interestingly, 17 HDAC3-regulated transcription factors, *Nfkb2*, *Trp53* and *Relb* transcription factors were induced by MHCII/TCR interactions with CD4^+^ thymocytes **(Figure 2I)**. Moreover, the expression of several cytokines, chemokines and cell adhesion molecules was also upregulated **(Figure 2J,K).** Altogether, these data show that CD4^+^ thymocytes control, through MHCII/TCR interactions, the functional properties of the fraction of mTEC^lo^ that express intermediate levels of MHCII and activate key transcriptional programs governing mTEC differentiation and function.

### MHCII/TCR interactions with CD4^+^ thymocytes regulate in mTEC^lo^ the development of Fezf2^+^ pre-Aire, post-Aire and tuft-like mTEC subsets

Since key transcription factors implicated in mTEC differentiation were upregulated in mTEC^lo^ by MHCII/TCR-mediated interactions with CD4^+^ thymocytes **(Figure 1G and 2I)**, we next analyzed the mTEC composition of ΔCD4 and mTEC^ΔMHCII^ mice. An Aire/Fezf2 co-staining both by histology and flow cytometry revealed a substantial reduction in Aire^-^Fezf2^+^ and Aire^+^Fezf2^+^ cells **(Figure 3A,B)**. We further analyzed by flow cytometry Aire and Fezf2 expression in mTEC^lo^ and mTEC^hi^ according to the gating strategy shown in **Figure S1A**. In agreement with the detection of *Aire* in the proliferating and maturational single-cell clusters in mTEC^lo^ (Baran-Gale et al, 2020; Dhalla et al, 2020), we found that Aire protein was expressed in a small fraction of mTEC^lo^ compared to mTEC^hi^ in WT, ΔCD4 and mTEC^ΔMHCII^ mice **(Figure 3B)**. Aire^-^ Fezf2^+^ and Aire^+^Fezf2^+^ mTECs were reduced in mTEC^lo^ of ΔCD4 and mTEC^ΔMHCII^ mice with a more marked effect in mTEC^hi^. This decrease was not due to impaired proliferation since normal frequencies of Ki-67^+^ proliferating of these cells were observed in ΔCD4 and mTEC^ΔMHCII^ mice **(Figure S4).** Furthermore, numbers of involucrin^+^TPA^+^Aire^-^ post-Aire cells were reduced in the medulla of ΔCD4 and mTEC^ΔMHCII^ mice **(Figure S5)**, consistently with the decrease of their Aire^+^ mTEC^hi^-precursors **(Figure 3B-C)**.

**Figure 3.**
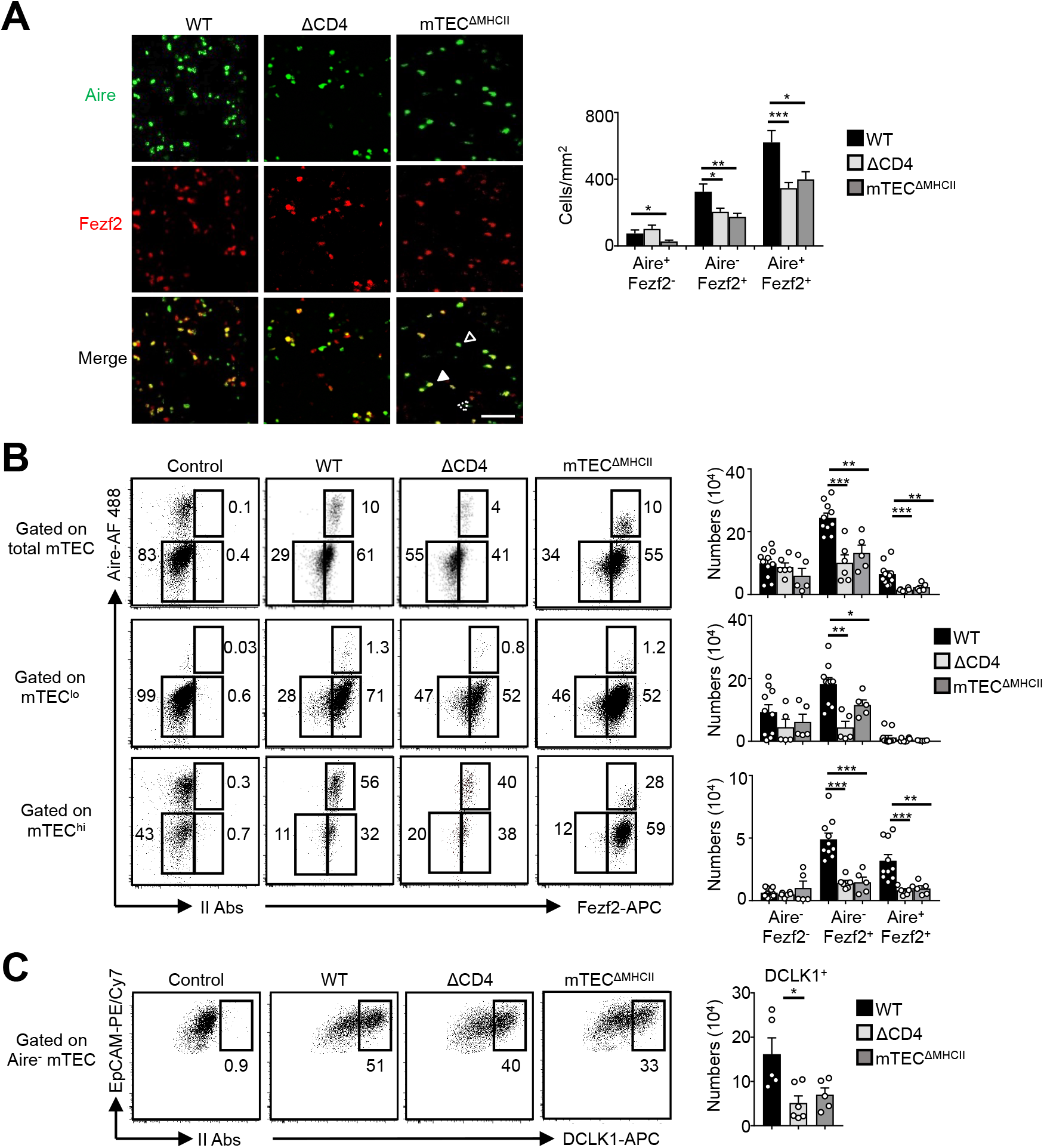
mTEC^lo^ subset composition is altered in ΔCD4 and mTEC^ΔMHCII^ mice. **(A)** Confocal images of thymic sections from WT, ΔCD4 and mTEC^ΔMHCII^ mice stained for Aire (green) and Fezf2 (red). 12 and 20 sections derived from 2 WT, 2 ΔCD4 and 2 mTEC^ΔMHCII^ mice were quantified. Scale bar, 50 μm. Unfilled, dashed and solid arrowheads indicate Aire^+^Fezf2^-^, Aire^-^Fezf2^+^ and Aire^+^Fezf2^+^ cells, respectively. The histogram shows the density of Aire^+^Fezf2^-^, Aire^-^Fezf2^+^ and Aire^+^Fezf2^+^ cells. **(B,C)** Flow cytometry profiles and numbers of Aire^-^Fezf2^-^, Aire^-^Fezf2^+^ and Aire^+^Fezf2^+^ cells in total mTECs, mTEC^lo^ and mTEC^hi^ **(B)** and of DCKL1^+^ cells in Aire^-^ mTECs **(C)** from WT, ΔCD4 and mTEC^ΔMHCII^ mice. II Abs: secondary antibodies. Data are representative of 2-3 independent experiments (n=3-4 mice per group and experiment). Error bars show mean◻±◻SEM, *p<0.05, **p<0.01, ***p<0.001, ****p<0.0001 using unpaired Student’s t-test for **A** and two-tailed Mann-Whitney test for **B, C**.

We also analyzed tuft-like mTECs since the expression of the transcription factor Pou2f3, known to control the development of this cell type (Bornstein et al, 2018; Miller et al, 2018), was decreased in mTEC^lo^ of ΔCD4 and mTEC^ΔMHCII^ mice **(Figure 1G, 2I).** We found that numbers of tuft-like mTECs identified by flow cytometry using the DCLK1 marker were also reduced in both mice **(Figure 3C)**, indicating that their development is controlled by MHCII/TCR interactions with CD4^+^ thymocytes. Importantly, Aire^-^Fezf2^+^ and Aire^+^Fezf2^+^ mTEC^lo^ and mTEC^hi^ as well as DCLK1^+^ tuft-like mTECs were similarly reduced in MHCII^-/-^ mice, confirming that CD4^+^ thymocytes control the cellularity of these novel mTEC subsets **(Figure S3B,C).** Altogether, these data reveal that MHCII/TCR-mediated interactions with CD4^+^ thymocytes have a broad impact on mTEC^lo^ composition by controlling the cellularity of Fezf2^+^ pre-Aire mTECs, post-Aire and tuft-like mTECs.

### Highly self-reactive CD4^+^ thymocytes activate maturational programs in mTEC^lo^

We next assessed the impact of highly self-reactive interactions with CD4^+^ thymocytes in mTEC^lo^ using OTII-*Rag2*^-/-^ and RipmOVAxOTII-*Rag2*^-/-^ transgenic mice. Both models possess CD4^+^ thymocytes expressing an MHCII-restricted TCR specific for the chicken ovalbumin (OVA). The Rip-mOVA line expresses a membrane-bound OVA form specifically in mTECs and consequently high affinity interactions between OVA-expressing mTECs and OTII CD4^+^ thymocytes are only possible in RipmOVAxOTII-*Rag2*^-/-^ mice (Kurts et al, 1996). In mTEC^lo^, we expect that the Aire-dependent mOVA is expressed in a fraction of cells that express intermediate levels of CD80 and MHCII, since these cells have been shown to start expressing Aire. In contrast to total Erk1/2 MAPK, p38 MAPK, IKKα and p65, the non-classical NF-kB subunit RelB was increased in mTEC^lo^ at mRNA and protein levels in RipmOVAxOTII-*Rag2*^-/-^ compared to OTII-*Rag2*^-/-^ mice **(Figure 4A,B and Figure S6)**. The level of RelB phosphorylation was also higher **(Figure 4B)**, indicating that self-reactive CD4^+^ thymocytes activate the non-classical NF-κB pathway in mTEC^lo^.

**Figure 4.**
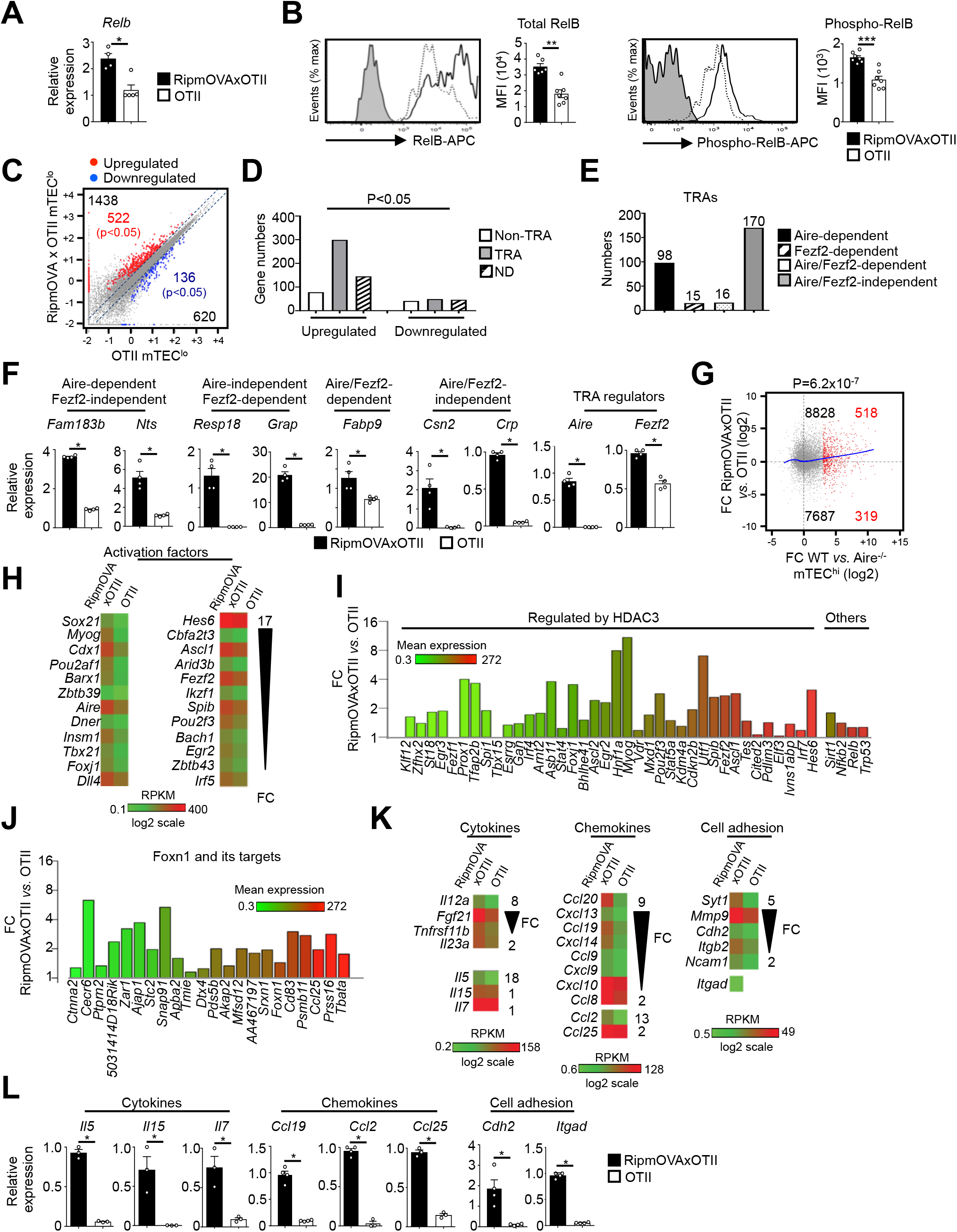
Highly self-reactive CD4^+^ thymocytes control the transcriptional and functional properties of mTEC^lo^. **(A)** *Relb* mRNA was measured by qPCR in mTEC^lo^ from RipmOVAxOTII-*Rag2*^-/-^ (n=4) and OTII-*Rag2*^-/-^ (n=5) mice. **(B)** Total and phospho-RelB (Ser552) were analyzed by flow cytometry in mTEC^lo^ from RipmOVAxOTII-*Rag2*^-/-^ and OTII-*Rag2*^-/-^ mice. Data are representative of 2 independent experiments (n=3-4 mice per group and experiment). **(C)** Scatter-plot of gene expression levels (FPKM) in mTEC^lo^ from RipmOVAxOTII-*Rag2*^-/-^ *versus* OTII-*Rag2*^-/-^ mice. Genes with fold difference ≥2 and p-adj <0.05 were considered as upregulated or downregulated genes (red and blue dots, respectively). RNA-seq was performed on 2 independent biological replicates with mTEC^lo^ derived from 5-8 mice. **(D)** Numbers of TRAs and non-TRAs in genes up- and downregulated in mTEC^lo^ from RipmOVAxOTII-*Rag2*^-/-^ *versus* OTII-*Rag2*^-/-^ mice. ND: not determined. **(E)** Numbers of induced Aire-dependent, Fezf2-dependent, Aire/Fezf2-dependent and Aire/Fezf2-independent TRAs. **(F)** Aire-dependent (*Fam183b, Nts*), Fezf2-dependent (*Resp18, Grap*), Aire/Fezf2-dependent (*Fabp9*), Aire/Fezf2-independent (*Csn2*, *Crp*) TRAs, *Aire* and *Fezf2* mRNAs were measured by qPCR in mTEC^lo^ from RipmOVAxOTII-*Rag2*^-/-^ (n=4) and OTII-*Rag2*^-/-^ (n=4) mice. **(G)** Scatter-plot of gene expression variation in mTEC^lo^ from RipmOVAxOTII-*Rag2*^-/-^ *versus* OTII-*Rag2*^-/-^ mice and in mTEC^hi^ from WT *versus* Aire^-/-^ mice. The loess fitted curve is shown in blue and induced Aire-dependent genes (FC>5) in red. **(H)** Heatmap of selected activation factors. **(I,J)** Expression fold change in HDAC3-induced transcriptional regulators and other transcription factors **(I)** and in Foxn1 targets **(J)** in mTEC^lo^ from RipmOVAxOTII-*Rag2*^-/-^ *versus* OTII-*Rag2*^-/-^ mice. The color code represents gene expression level. **(K)** Heatmap of selected cytokines, chemokines and cell adhesion molecules. **(L)** *Il5*, *Il15, Il7, Ccl19, Ccl2, Ccl25, Cdh2* and *Itgad* mRNAs were measured by qPCR in mTEC^lo^ from RipmOVAxOTII-*Rag2*^-/-^ (n=3-4) and OTII-*Rag2*^-/-^ (n=3-4) mice. Error bars show mean◻±◻SEM, *p<0.05, **p<0.01, ***p<0.001 using two-tailed Mann-Whitney test for **A, B, F, L** and Chi-squared test for **D, G**.

To define the genome-wide effects of highly self-reactive CD4^+^ thymocytes in mTEC^lo^, we compared the gene expression profiles of mTEC^lo^ from RipmOVAxOTII-*Rag2*^-/-^ versus OTII-*Rag2*^-/-^ mice **(Figure S1B)** and found an upregulation of 1438 genes (FC>2) reaching statistical significance for 522 of them (Cuffdiff p<0.05). 620 genes were also downregulated (FC<0.5) with 136 reaching significance (Cuffdiff p<0.05) **(Figure 4C)**. The genes upregulated exhibited a ~4-fold more of TRA over non-TRA genes (p=4.7×10^-23^), which corresponded mainly to Aire-dependent and Aire/Fezf2-independent TRAs **(Figure 4D-F and Table S3)**. Similarly to the WT versus mTEC^ΔMHCII^ comparison, we found a strong correlation (p=6.2×10^-7^) between the genes upregulated by self-reactive CD4^+^ thymocytes and the responsiveness of genes to Aire’s action obtained from the comparison between WT and *Aire*^-/-^ mTEC^hi^ **(Figure 4G)**. These results support an impact of antigen-specific interactions in the expression of TRAs in mTEC^lo^, notably on Aire-dependent TRAs. Importantly, these results are in agreement with the induction of a list of activation factors including *Aire* and *Fezf2* among of the non-TRA genes **(Figure 4H,I).** Similarly to the comparisons of the WT versus ΔCD4 or mTEC^ΔMHCII^, numerous HDAC3-induced regulators as well as *Sirt1*, *Nfkb2*, *Relb* and *Trp53* transcription factors were upregulated in mTEC^lo^ of RipmOVAxOTII-*Rag2*^-/-^ mice **(Figure 4J).** Interestingly, the Foxn1 transcription factor, implicated in TEC differentiation and growth, 21 out of its top 30 targets (Zuklys et al, 2016), cytokines, chemokines and cell adhesion molecules were also upregulated **(Figure 4K-L).** Altogether, these data reveal that highly self-reactive CD4^+^ thymocytes control in the mTEC^lo^ fraction that express intermediate levels of MHCII not only key transcription factors driven their differentiation but also crucial molecules for T-cell development and selection such as TRAs, cytokines, chemokines and adhesion molecules.

### Highly self-reactive CD4^+^ thymocytes control mTEC subset composition from a progenitor stage

Given that highly self-reactive CD4^+^ thymocytes induce key transcription factors in mTEC^lo^ (Figure 4H, I), we examined their respective impact on mTEC subset development. Strikingly, numbers of total TECs and mTECs were higher in RipmOVAxOTII-*Rag2*^-/-^than in OTII-*Rag2*^-/-^mice (Figure 5A,B).We analyzed four TEC subsets based on MHCII and UEA-1 levels, as previously described (Lopes et al, 2017; Wong et al, 2014) (Figure 5C).In contrast to cTEChi (MHCIIhiUEA-1lo), numbers of TEClo (MHCIIloUEA-1lo), mTEC^lo^ (MHCIIloUEA-1+) and mTEChi (MHCIIhiUEA-1+) were higher in RipmOVAxOTII-*Rag2*^-/-^than in OTII-*Rag2*^-/-^mice. Interestingly, numbers of α6-integrinhiSca-1hi thymic epithelial progenitor (TEPC)-enriched cells in the TEClo subset were also increased (Figure 5D),indicating that self-reactive CD4^+^ thymocytes control TEC development from a precursor stage. Of note, this strategy of TEC identification was not possible in ΔCD4 and mTECΔMHCII mice since MHCII expression is abrogated in TECs of these mice (Irla et al, 2008).

**Figure 5.**
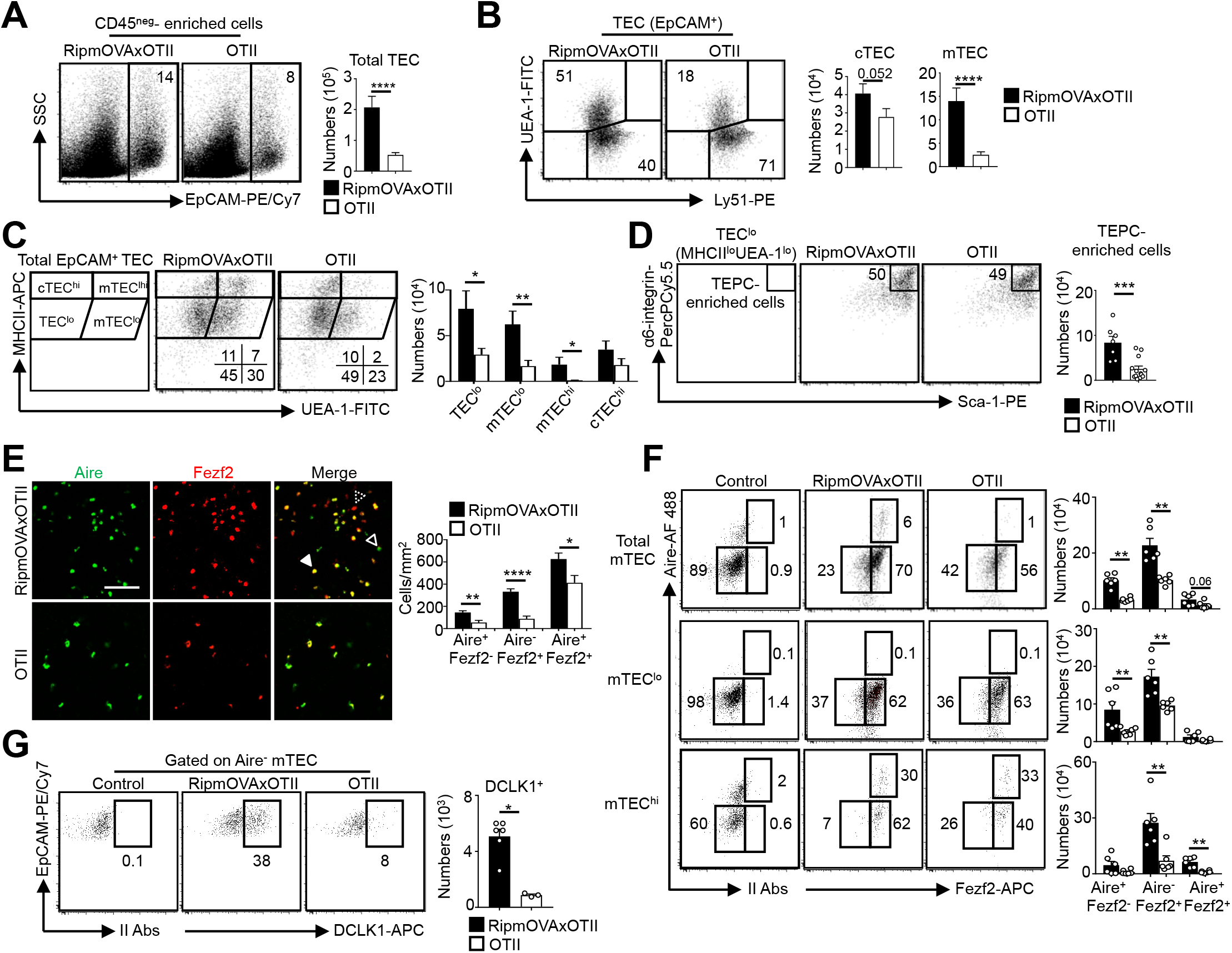
Highly self-reactive CD4^+^ thymocytes control mTEC development from an early progenitor stage. **(A-D)** Flow cytometry profiles and numbers of total TECs (EpCAM^+^) **(A)**, cTECs (UEA-1^-^Ly51^hi^), mTECs (UEA-1^-^Ly51^lo^) **(B)**, TEC^lo^ (MHCII^lo^UEA-1^lo^), cTEC^hi^ (MHCII^hi^UEA-1^lo^), mTEC^lo^ (MHCII^lo^UEA-1^hi^) and mTEC^hi^ (MHCII^hi^UEA-1^hi^) **(C)**, α6-integrin^hi^Sca-1^hi^ TEPC-enriched cells in TEC^lo^ **(D)** in CD45^neg^-enriched cells from RipmOVAxOTII-*Rag2*^-/-^ and OTII-*Rag2*^-/-^ mice. Data are representative of 4 experiments (n=3 mice per group and experiment). **(E)** Confocal images of thymic sections from RipmOVAxOTII-*Rag2*^-/-^ and OTII-*Rag2*^-/-^ mice stained for Aire (green) and Fezf2 (red). 11 and 22 sections derived from 2 RipmOVAxOTII-*Rag2*^-/-^ and OTII-*Rag2*^-/-^ mice were quantified, respectively. Scale bar, 50 μm. Unfilled, dashed and solid arrowheads indicate Aire^+^Fezf2^lo^, Aire^-^Fezf2^+^ and Aire^+^Fezf2^+^ cells, respectively. The histogram shows the density of Aire^+^Fezf2^lo^, Aire^-^Fezf2^+^ and Aire^+^Fezf2^+^ cells. **(F,G)** Flow cytometry profiles and numbers of Aire^-^Fezf2^-^, Aire^-^Fezf2^+^ and Aire^+^Fezf2^+^ cells in total mTECs, mTEC^lo^ and mTEC^hi^ **(F)** and of DCKL1^+^ cells in Aire^-^ mTECs **(G)** from RipmOVAxOTII-*Rag2*^-/-^ and OTII-*Rag2*^-/-^ mice. II Abs: secondary antibodies. Data are representative of 2 independent experiments (n=3-4 mice per group and experiment). Error bars show mean◻±◻SEM, *p<0.05, **p<0.01, ***p<0.001, ****p<0.0001 using unpaired Student’s t-test for **A-E** and two-tailed Mann-Whitney test for **F,G**.

A higher density of Aire^+^Fezf2^-^, Aire^-^Fezf2^+^ and Aire^+^Fezf2^+^ cells was observed in medullary regions of RipmOVAxOTII-*Rag2*^-/-^ mice by immunohistochemistry **(Figure 5E).** Furthermore, numbers of Aire^-^Fezf2^-^ mTEC^lo^ analyzed by flow cytometry were also higher in RipmOVAxOTII-*Rag2*^-/-^ mice, confirming that self-reactive CD4^+^ thymocytes control mTEC differentiation from an early stage **(Figure 5F)**. Numbers of Aire^-^Fezf2^+^ and Aire^+^Fezf2^+^ mTEC^lo^ and mTEC^hi^ were also markedly increased in these mice, although similar frequencies of proliferating Ki-67^+^ cells was observed **(Figure S7).** In agreement with increased Aire^+^ mTECs, involucrin^+^TPA^+^Aire^-^ post-Aire cells were enhanced **(Figure S8).** Altogether, these results demonstrate that highly self-reactive CD4^+^ thymocytes regulate mTECs from an early to a late developmental stage and thus the heterogeneity of mTECs in the thymus.

### Self-reactive CD4^+^ thymocytes enhance the level of active H3K4me3 mark in mTEC^lo^

Since histone modifications constitute important regulatory mechanisms that control the open and closed state of mTEC chromatin (Ucar & Rattay, 2015), we investigated whether self-reactive CD4^+^ thymocytes induce histone modifications in mTECs. We first analyzed in WT mTEC^lo^ the repressive H3K27me3 and the active H3K4me3 marks using chromatin immunoprecipitation (ChIP) followed by high-throughput sequencing (ChIP-seq). As expected, metagene analyses showed that Aire-dependent TRAs had higher levels of H3K27me3 in their genes than in all genes of the genome, confirming that they are in a repressive state **(Figure 6A)**. In contrast, Fezf2-dependent TRAs had reduced H3K27me3 levels and a significant enrichment of H3K4me3 in their transcriptional start site (TSS) **(Figure 6B)**. Similarly, Aire/Fezf2-independent TRAs were associated with low levels of H3K27me3 in their genes and high levels of H3K4me3 in their TSS. Thus, in contrast to Aire-dependent TRAs that are associated with the repressive H3K27me3 histone mark, Fezf2-dependent and Aire/Fezf2-independent TRAs are associated with the active H3K4me3 mark, indicating that these distinct TRAs are subjected to a distinct epigenetic regulation.

**Figure 6.**
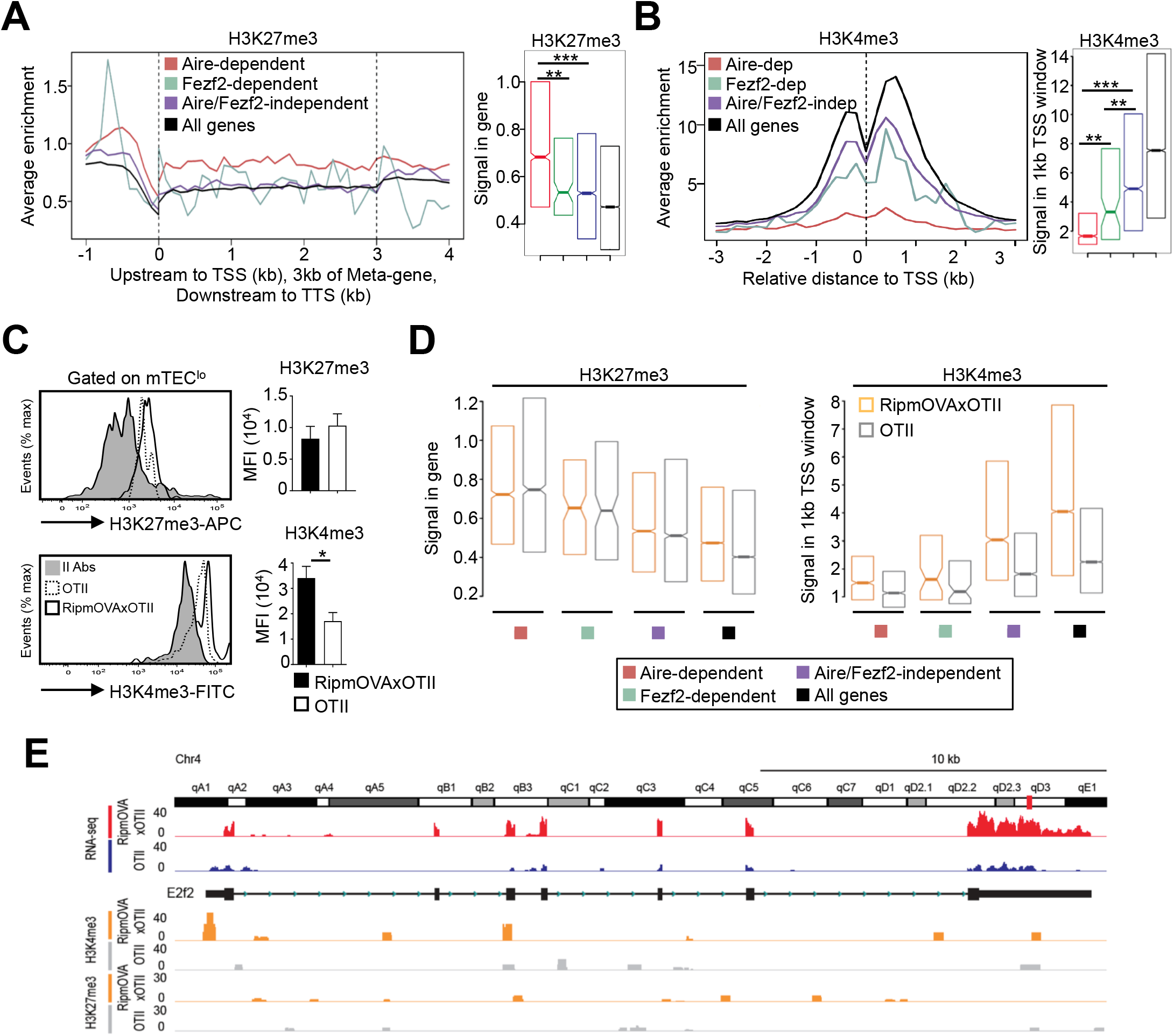
H3K27me3 and H3K4me3 landscape in TRA genes of mTEC^lo^ from WT, RipmOVAxOTII-*Rag2*^-/-^ and OTII-^Rag2-/-^ mice. **(A-B)** Metagene profiles of the average normalized enrichment of H3K27me3 **(A)** and H3K4me3 **(B)** against input for Aire-dependent, Fezf2-dependent and Aire/Fezf2-independent TRAs as well as for all genes of WT mTEC^lo^. Boxplots represent the median enrichment, the 95% CI of the median (notches) and the 75^th^ and 25^th^ percentiles of H3K27me3 and H3K4me3. **(C)** H3K27me3 and H3K4me3 levels were analyzed by flow cytometry in mTEC^lo^ from RipmOVAxOTII-*Rag2*^-/-^ and OTII-*Rag2*^-/-^ mice. Histograms show the MFI. **(D)** Boxplots represent the median enrichment of H3K27me3 and H3K4me3 of Aire-dependent, Fezf2-dependent and Aire/Fezf2-independent TRAs and in all genes of mTEC^lo^ from RipmOVAxOTII-*Rag2*^-/-^ and OTII-*Rag2*^-/-^ mice. **(E)** Expression (RNA-seq) and H3K27me3 and H3K4me3 chromatin state (ChIP-seq) of the Aire/Fezf2-independent TRA, *E2f2*, in mTEC^lo^ from RipmOVAxOTII-*Rag2*^-/-^ and OTII-*Rag2*^-/-^ mice. *p<0.05, **p<10^-3^, ***p<10^-7^, using the Mann-Whitney test for **A** and **B** and unpaired Student’s t-test for **C**.

We next assessed whether self-reactive CD4^+^ thymocytes control the H3K27me3 and H3K4me3 chromatin landscape in mTEC^lo^. In contrast to H3K27me3, we found an increased global level of H3K4me3 in RipmOVAxOTII-*Rag2*^-/-^ compared to OTII-*Rag2*^-/-^ mice by flow cytometry **(Figure 6C)**. We further analyzed by nano-ChIP-seq whether self-reactive CD4^+^ thymocytes regulate in mTEC^lo^ the level of these two histone marks in Aire-dependent, Fezf2-dependent and Aire/Fezf2-independent TRA genes. H3K27me3 levels in Aire-dependent TRA genes were comparable in mTEC^lo^ from RipmOVAxOTII-*Rag2*^-/-^ and OTII-*Rag2*^-/-^ mice **(Figure 6D; left panel)**. Although lower, H3K27me3 levels in Fezf2-dependent and Aire/Fezf2-independent TRAs as well as in all genes were similar in both mice, indicating that the interactions with self-reactive CD4^+^ thymocytes do not regulate this repressive mark in TRA genes **(Figure 6D; left panel).** In contrast, H3K4me3 global level was increased in the TSS of all TRAs in RipmOVAxOTII-*Rag2*^-/-^ compared to OTII-*Rag2*^-/-^ mice as well as in all genes **(Figure 6D; right panel)**. For representation, whereas the Aire/Fezf2-independent TRA, E2F transcription factor 2 (*E2f2*) induced by these interactions, was barely devoid of H3K27me3 in both mice, it was marked by H3K4me3 in its TSS specifically in RipmOVAxOTII-*Rag2*^-/-^ mice **(Figure 6E)**. These results thus show that self-reactive CD4^+^ thymocytes enhance the global level of the active H3K4me3 histone mark in mTEC^lo^ and in particular in the TSS of Fezf2-dependent and Aire/Fezf2-independent TRAs, indicative of an epigenetic regulation for their expression.

### MHCII/TCR interactions between mTECs and CD4^+^ thymocytes prevent the development of autoimmunity

We next evaluated the impact of mTEC-CD4^+^ thymocyte interactions on the generation of a self-tolerant T-cell repertoire by taking advantage that mTEC^ΔMHCII^ mice, in which MHCII/TCR interactions between mTECs and CD4^+^ thymocytes are abrogated, develop CD4^+^ and CD8^+^ T cells. Interestingly, since TRAs induced by MHCII/TCR interactions showed a diverse peripheral tissue distribution in mTEC^lo^ **(Figure 7A and Table S4)**, we analyzed by flow cytometry whether the TCRVβ repertoire is altered in mTEC^ΔMHCII^ mice. The TCRVβ usage was more altered in CD69^-^ mature CD4^+^ thymocytes than in CD69^-^ mature CD8^+^ thymocytes **(Figure 7B).** Some TCRVβ specificities were also altered in splenic CD4^+^ and CD8^+^ T cells. To determine whether these T cells contain self-reactive specificities, we adoptively transferred splenocytes from mTEC^ΔMHCII^ or WT mice into lymphopenic *Rag2*^-/-^recipients **(Figure 7C).** Mice that received splenocytes derived from mTEC^ΔMHCII^ mice lost significantly more weight than mice transferred with WT splenocytes **(Figure 7D).** They also exhibited splenomegaly with increased follicle areas and T-cell numbers showing a CD62L^lo^CD44^hi^ effector and CD62L^hi^CD44^hi^ central memory phenotype **(Figure 7E-G)**. Immune infiltrates in lungs and salivary glands were observed by histology and flow cytometry in 75% and 41% of mice, respectively **(Figure 7H,I)**. These two tissues contained increased numbers of central memory as well as CD44^+^CD69^+^ and CD44^+^CD69^-^ activated CD4^+^ and CD8^+^ T cells **(Figure 7J).** T-cell infiltrates were also observed in other tissues such as kidney, liver and colon **(Figure 7K).** Altogether, these data show that in absence of MHCII/TCR interactions between mTECs and CD4^+^ thymocytes, T cells contain self-reactive specificities and thus that these interactions are critical to the establishment of T-cell tolerance.

**Figure 7.**
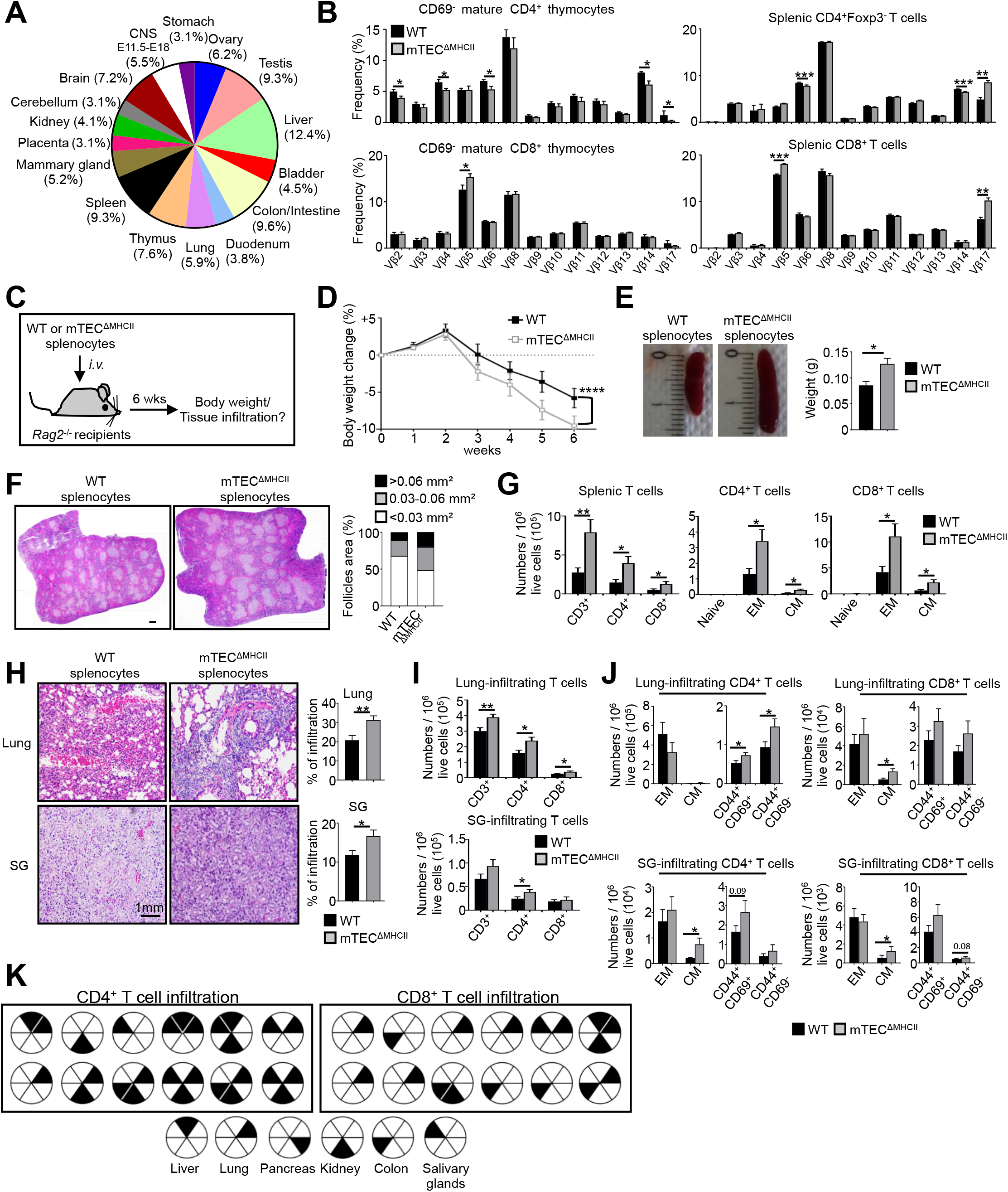
The adoptive transfer of splenocytes from mTEC ^ΔMHCII^ mice in *Rag2*^-/-^ recipients induces autoimmunity. **(A)** TRAs underexpressed in mTEC^lo^ from mTEC^ΔMHCII^ mice were assigned to their peripheral expression. **(B)** TCRVβ usage by CD69^-^ mature CD4^+^ and CD8^+^ thymocytes (left panel) and CD4^+^Foxp3^-^ and CD8^+^ splenic T cells (right panel) from WT and mTEC^ΔMHCII^ mice. **(C)** Body weight of *Rag2*^-/-^ mice transferred with WT or mTEC^ΔMHCII^ splenocytes was monitored during 6 weeks and tissue infiltration was examined. **(D)** Weight loss relative to the initial weight. **(E,F)** Representative spleen pictures and their weights **(E)** and hematoxylin/eosin counterstained splenic sections **(F)**. Scale bar, 1 mm. The histogram shows follicle areas. **(G)** Numbers of splenic CD3^+^, CD4^+^ and CD8^+^ T cells and of naive (CD44^lo^CD62L^hi^), effector memory (EM; CD44^hi^CD62L^lo^) and central memory (CM; CD44^hi^CD62L^hi^) phenotype. **(H)** Lung and salivary gland (SG) immune infiltrates detected by hematoxylin/eosin counterstaining. Scale bar, 1 mm. **(I,J)** Numbers of T cells **(I)** and of naive, effector and central memory phenotype as well as CD44^+^CD69^+^ and CD44^+^CD69^-^ T cells **(J)** in lungs and SG. **(K)** Schematic of T-cell infiltrates in mice transferred with mTEC^ΔMHCII^ splenocytes relative to those transferred with WT splenocytes. Each circle and black triangles represent an individual mouse and T-cell infiltration in a specific tissue, respectively. Data are representative of 2 independent experiments (n=5-7 mice per group and experiment). Error bars show mean◻±◻SEM, ****p<0.0001 using two-way ANOVA for **D** and unpaired Student’s t-test for **B** and **E-J**. *p<◻0.05, **p<0.01, ***p<0.001.

## Discussion

Since mTECs play a crucial role in immunological tolerance by expressing TRAs, it is essential to deepen our knowledge of the mechanisms that sustain their development. Here using three distinct transgenic models, we found that self-reactive CD4^+^ thymocytes control the developmental transcriptional programs from the mTEC^lo^ stage. CD4^+^ thymocytes induce in mTEC^lo^ the phosphorylation of p38 MAPK and IKKα, the latter implicated in mTEC development (Lomada et al, 2007; Shen et al, 2019). Moreover, self-reactive CD4^+^ thymocytes induce RelB expression and increase its phosphorylation level. Interestingly, this non-classical NF-κB subunit is crucial for mTEC development and Aire-dependent and independent TRA expression (Riemann et al, 2017). These data thus show that CD4^+^ thymocyte development and antigen-specific interactions play non-redundant roles in activating intracellular pathways from the mTEC^lo^ stage. Analysis of the mTEC^lo^ transcriptional landscape by high throughput RNA-seq revealed that self-reactive CD4^+^ thymocytes also induce *Nfkb2* (p52), known to form an heterocomplex with RelB in the nucleus upon activation (Irla et al, 2010). p52 is important for mTEC development, Aire and TRA expression (Zhang et al, 2006; Zhu et al, 2006). Consequently, ΔCD4, mTEC^ΔMHCII^ and OTII-*Rag2*^-/-^ mice in which MHCII/TCR interactions between mTECs and CD4^+^ thymocytes are disrupted have altered *Relb* and *Nfkb2* expression, reduced Aire^+^ mTEC numbers and Aire-dependent TRA representation. Our results are in agreement with the fact that RANK signaling is activated by membrane-bound RANKL and not soluble RANKL and thereby in the context of physical interactions between mTECs and CD4^+^ thymocytes (Asano et al, 2019).

These interactions also induce *Trp53* (p53) that controls the mTEC niche (Rodrigues et al, 2017) and *Irf4* and *Irf7* transcription factors that regulate key chemokines implicated in thymocyte medullary localization and mTEC differentiation (Haljasorg et al, 2017; Otero et al, 2013). Furthermore, the deacetylase Sirtuin-1 (*Sirt1*), which regulates Aire activity (Chuprin et al, 2015) and *Spib* that limits mTEC differentiation (Akiyama et al, 2014) were also induced. Self-reactive CD4^+^ thymocytes thus induce key transcription factors that not only positively but also negatively control mTEC differentiation. Remarkably, our three different transgenic models revealed that CD4^+^ thymocytes induce HDAC3-dependent mTEC-specific transcription factors (Goldfarb et al, 2016). Among them, *Pou2f3* is involved in tuft-like mTEC development (Bornstein et al, 2018; Miller et al, 2018), which is consistent with our results that self-reactive CD4^+^ thymocytes control the cellularity of these cells. Our data thus identify that CD4^+^ thymocytes control in mTEC^lo^ the expression of master transcriptional regulators of mTEC differentiation and function.

In line with these results, we found that self-reactive CD4^+^ thymocytes control the differentiation of distinct mTEC subsets. Interestingly, self-reactive CD4^+^ thymocytes regulate TEC development from a precursor stage since they increase numbers of TEPC-enriched cells that express non-negligible MHCII levels. We provide the first evidence that they control Fezf2^+^ pre-Aire mTECs. Accordingly, the expression of Fezf2 and its respective TRAs was enhanced by CD4^+^ thymocytes. Moreover, self-reactive CD4^+^ thymocytes regulate tuft-like and post-Aire cell numbers in mTEC^lo^. These results thus reveal that antigen-specific interactions with CD4^+^ thymocytes have an unsuspected broad impact on mTEC composition by driving their development from an early progenitor to a late post-Aire stage.

Interestingly, high-throughput RNA-seq showed that MHCII/TCR interactions with CD4^+^ thymocytes upregulate the expression of chemokines in mTECs. Among them, CCL19 (CCR7 ligand) is implicated in the medulla localization of thymocytes and the emigration of newly generated T cells (Ueno et al, 2004); and CCL22 (CCR4 ligand) in medullary entry and thymocyte/dendritic cell interactions (Hu et al, 2015). Self-reactive CD4^+^ thymocytes also enhance CCL2 (CCR2 ligand) and CCL20 (CCR6 ligand) that promote the entry of peripheral dendritic cells and Foxp3^+^ regulatory T cells into the thymus (Baba et al, 2009; Cowan et al, 2018; Lopes et al, 2018). mTEC-CD4^+^ thymocyte interactions thus induce key chemokines that regulate the trafficking of thymocytes and thymic entry of peripheral cells that participate in tolerance induction. Moreover, cytokines such as *Il15* and *Fgf21* implicated in invariant NKT development and TEC protection against senescence, as well as adhesion molecules involved in mTEC-thymocyte interactions were also induced (Pezzi et al, 2016; White et al, 2014; Youm et al, 2016). Altogether, these results show that self-reactive CD4^+^ thymocytes regulate functional properties of mTEC^lo^ by inducing chemokines, cytokines and adhesion molecules that are critical for T-cell development.

The expression of TRAs is regulated by Aire and at a lesser extent by Fezf2 (Anderson et al, 2002; Takaba et al, 2015). In agreement with other studies (Gray et al, 2007; Takaba et al, 2015), we found Fezf2 in both mTEC^lo^ and mTEC^hi^ whereas Aire protein is mainly expressed in mTEC^hi^. Nevertheless and in line with recent single-cell transcriptomic analyses (Baran-Gale et al, 2020; Dhalla et al, 2020), we detected Aire by flow cytometry, qPCR and RNA-seq in a small subset (~1.5%) of mTEC^lo^. CD4^+^ thymocyte interactions upregulate *Aire* and *Fezf2* and some of their respective TRAs in these cells. Interestingly, in contrast to Aire-dependent TRAs that are characterized by high levels of H3K27me3 (Handel et al, 2018; Sansom et al, 2014), we found that Fezf2-dependent TRAs show high levels of H3K4me3. This highlights that Aire and Fezf2 use distinct epigenetic modes in regulating TRA expression. Remarkably, these interactions also induce in mTEC^lo^ numerous Aire/Fezf2-independent TRAs, whose regulation remains unknown. Similarly to Fezf2-dependent TRAs, they had high levels of H3K4me3 in their TSS, suggesting that Aire/Fezf2-independent TRAs are not subjected to the same regulatory transcriptional mechanisms than Aire-dependent TRAs. Our results are consistent with a previous study indicating that the Aire-independent TRA, *Gad67*, shows active epigenetic marks (Tykocinski et al, 2010). Remarkably, self-reactive CD4^+^ thymocytes increase H3K4me3 level in the TSS of all TRA categories, thus providing a novel epigenetic mechanistic insight into how they regulate the mTEC gene expression profile. In line with TRA regulation and the development of distinct mTEC subsets, the TCRVβ repertoire of mature T cells was altered when MHCII/TCR interactions were abrogated between mTECs and CD4^+^ thymocytes. Importantly, mTEC-CD4^+^ thymocyte interactions are critical for the generation of a self-tolerant T-cell repertoire since the adoptive transfer of splenocytes from mTEC^ΔMHCII^ mice into *Rag2*^-/-^ recipients led to signs of autoimmunity.

In summary, our genome-wide scale study reveals that self-reactive CD4^+^ thymocytes activate in mTEC^lo^ transcriptional programs that sustain the differentiation into Aire^+^ mTEC^hi^-precursors, Aire^+^Fezf2^+^, post-Aire and tuft-like mTECs. These interactions also induce TRAs, cytokines, chemokines and adhesion molecules that are all implicated in mTEC function. Thus, CD4^+^ thymocytes control several unsuspected aspects of mTEC^lo^ required for the establishment of T-cell tolerance.

## Acknowledgments

We gratefully thank Pr. Walter Reith for *Ciita^III+IV^*^-/-^ and K14x*Ciita^III+IV^*^-/-^ mice and Dr. Bruno Lucas for MHCII^-/-^ mice. We also thank Lionel Chasson for help with paraffin-embedded tissues. We acknowledge the flow cytometry and animal facility platforms at CIML for excellent technical support. This project was funded by the Marie Curie Actions (Career Integration Grants, CIG_SIGnEPI4Tol_618541 to M.I.), the Agence Nationale de la Recherche (2011-CHEX-001-R12004KK to M.G), the ‘Fondation Princesse Grace de la Principauté de Monaco (to P.F) and institutional grants from Institut National de la Santé et de la Recherche Médicale, Centre National de la Recherche Scientifique and Aix-Marseille Université. We acknowledge financial support from n°ANR-10-INBS-04-01 France Bio Imaging and the France Génomique national infrastructure, funded as part of the “Investissements d’Avenir” program managed by the Agence Nationale de la Recherche (ANR-10-INBS-0009). N.L. and J.C. are supported by a PhD fellowship from Aix-Marseille University and the Ministère de l’Enseignement Supérieur et de la Recherche, respectively.

## Author contributions

N.L., N.B., J.C. and M.I. performed experiments and analyzed the data. M.G. performed all the bioinformatics analysis and interpreted the data. P.F. participated to the conceptualization of ChIP-seq experiments. N.L and M.I. wrote the manuscript. M.I. initiated, supervised and conceived the study.

## Materials and Methods

### Mice

C57BL/6 WT mice were purchased from Charles River. *Ciita^III+IV^*^-/-^ (ΔCD4) (LeibundGut-Landmann et al, 2004), MHCII^-/-^ (Madsen et al, 1999), K14x*Ciita^III+IV^*^-/-^ (mTEC^ΔMHCII^) (Irla et al, 2008), OTII (Barnden et al, 1998), RipmOVAxOTII (Kurts et al, 1996) and *Rag2*^-/-^ (Shinkai et al, 1992) mice were on C57BL/6J background. OTII and RipmOVAxOTII were backcrossed on *Rag2*^-/-^ background. All mice were maintained under specific pathogen-free conditions at an ambient temperature of 22°C at the animal facilities of the CIML (Marseille, France). Standard food and water were given *ad libitum*. Males and females were used at the age of 5-6 weeks. Experiments were performed in accordance with the animal care guidelines of the European Union and French laws. All experiments were done in accordance with national and European laws for laboratory animal welfare (EEC Council Directive 2010/63/UE), and were approved by the Marseille Ethical Committee for Animal Experimentation (Comité National de Réflexion Ethique sur l’Expérimentation Animale no. 14).

### mTEC purification

mTECs were isolated by enzymatic digestion with 50 μg/ml of liberase TM (Roche) and 100 μg/ml of DNase I (Roche) in HBSS medium, as previously described (Lopes et al, 2017). CD45^+^ hematopoietic cells were depleted using anti-CD45 magnetic beads by autoMACS with the depleteS program (Miltenyi Biotec). Total mTECs (EpCAM^+^UEA-1^+^Ly51^lo^), mTEC^lo^ (EpCAM^+^UEA-1^+^Ly51^lo^CD80^lo/int^) and mTEC^hi^ (EpCAM^+^UEA-1^+^Ly51^lo^CD80^hi^) were sorted with a FACSAriaIII cell sorter (BD). The purity of sorted mTEC^lo^ was >98%. Flow cytometry gating strategies are shown in **Figure S1**.

### mTEC antigen presentation assays

Variable numbers of mTECs from WT and mTEC^ΔMHCII^ mice loaded or not with OVA_323-339_ (5μM, PolyPeptide group) were co-cultured with 10^5^ OTII CD4^+^ T cells (purified with a CD4^+^ T cell isolation kit, Miltenyi Biotec) in RPMI medium (ThermoFisher) supplemented with 10% FCS (Sigma Aldrich), L-glutamine (2◻mM, ThermoFisher), sodium pyruvate (1◻mM, ThermoFisher), 2-mercaptoethanol (2◻×◻10^-5^ M, ThermoFisher), penicillin (100◻IU◻per ml, ThermoFisher) and streptomycin (100◻μg◻per ml, ThermoFisher). The activation of OTII CD4^+^ T cells was assessed 18h later by flow cytometry based on the upregulation of the CD69 marker.

### Flow Cytometry

TECs, thymocytes and splenic T cells were analyzed by flow cytometry (FACS Canto II, BD) with standard procedures. Cells were incubated for 15 min at 4°C with Fc-block (anti-CD16/CD32, 2.4G2, BD biosciences) before staining. Antibodies are listed in **Table S5**. For intracellular staining with anti-Foxp3, anti-Ki-67, anti-p38 MAPK, anti-phospho p38 MAPK (Thr180/Tyr182), anti-IKKα, anti-phospho IKKα(Ser180)/IKKβ(Ser181), anti-Erk1/2 MAPK, anti-phospho Erk1/2 MAPK (Thr202/Tyr204), anti-p65, anti-phospho p65(ser536), anti-RelB, anti-phospho RelB(ser552) and anti-DCLK1 antibodies, cells were fixed, permeabilized and stained with the Foxp3 staining kit according to the manufacturer’s instructions (eBioscience). Intracellular staining with anti-Aire, anti-Fezf2, anti-H3K4me3 and anti-H3K27me3 antibodies was performed with Fixation/Permeabilization Solution Kit (BD). Secondary antibodies (II Abs) were used to set positive staining gates. Flow cytometry analysis was performed with a FACSCanto II (BD) and data were analyzed using FlowJo software (BD).

### Quantitative RT-PCR

Total RNA was prepared with TRIzol (Invitrogen). cDNAs was synthesized with oligo(dT) using Superscript II reverse transcriptase (Invitrogen). qPCR was performed with the ABI 7500 fast real-time PCR system (Applied Biosystems) and SYBR Premix Ex Taq master mix (Takara). Primers are listed in **Table S6**.

### *In vivo* transfer of splenocytes into *Rag2*^-/-^ recipients

3.10^6^ splenocytes from WT and mTEC^ΔMHCII^ mice of 8 weeks of age were intravenously injected into *Rag2^-/-^* female recipients. CD3^+^, CD4^+^ and CD8^+^ T-cell infiltrates were analyzed 6 weeks after transfer by histology and flow cytometry in different peripheral tissues.

### Histology

Tissues were fixed in 10% buffered formalin (Sigma) and embedded in paraffin blocks. 4μm thick sections were stained with hematoxylin-eosin (Thermofisher) and analyzed by light microscopy (Nikon Statif eclipse Ci-L). For immunofluorescence experiments, frozen thymic sections were stained as previously described(Serge et al, 2015) with Alexa Fluor 488 or Alexa Fluor 647-conjugated anti-Aire (5H12; eBioscience), biotinylated anti-TPA (LTL; Vector Laboratories), rabbit anti-Fezf2 (F441; IBL Tecan) and rabbit anti-involucrin (BioLegend) antibodies. Rabbit anti-Fezf2 and rabbit anti-involucrin were revealed with Cy3-conjugated anti-rabbit IgG (Invitrogen) and biotinylated anti-TPA was revealed with Alexa Fluor 488-conjugated streptavidin (Invitrogen). Sections were mounted with Mowiol (Calbiochem). Immunofluorescence confocal microscopy was performed with a LSM780 Leica SP5X confocal microscope. Images were analyzed with ImageJ Software.

### RNA-seq experiments

Total RNA purified from mTEC^lo^ **(Figure S1)** was extracted with miRNeasy® Micro Kit (Qiagen) and RNA quality was assessed on an Agilent 2100 BioAnalyzer (Agilent Technologies). RNA Integrity Number values over 8 were obtained. RNA-seq libraries were generated using the SMART-Seq-v4-Ultra Low Input RNA Kit (Clontech) combined to the Nextera library preparation kit (Illumina) following manufacturers’ instructions. Libraries were sequenced with the Illumina NextSeq 500 machine to generate datasets of single-end 75bp reads. Two independent biological replicates were used per each condition. RNA-seq analysis is detailed in the Supplemental Methods.

### Nano-ChIP-seq experiments

Nano-ChIP-seq was performed as previously described (Adli & Bernstein, 2011) on 5.10^4^ purified mTEC^lo^ **(Figure S1)**. ChIP-seq libraries were prepared with TruSeq ChIP Sample Preparation Kit (Illumina) and 2×75-bp paired-end reads were sequenced on an Illumina HiSeq. ChIP-seq analysis is detailed in the Supplemental Methods.

### Statistics

Data are presented as means ± standard error of mean (SEM). Statistical analysis was performed with GraphPad Prism 7.03 software by using ANOVA, Chi-square, unpaired Student’s t test or Mann-Whitney test. ****, p<0.0001; ***, p<0.001; **, p<0.01; *, p<0.05. Normal distribution of the data was assessed using d’Agostino-Pearson omnibus normality test.

### Data availability

RNA-seq and ChIP-seq data reported in this paper were deposited with Gene Expression Omnibus under accession numbers: GSE144650 and GSE144680.

## References

Adli M, Bernstein BE (2011) Whole-genome chromatin profiling from limited numbers of cells using nano-ChIP-seq. Nat Protoc 6: 1656–1668

Akiyama N, Shinzawa M, Miyauchi M, Yanai H, Tateishi R, Shimo Y, Ohshima D, Matsuo K, Sasaki I, Hoshino K, Wu G, Yagi S, Inoue J, Kaisho T, Akiyama T (2014) Limitation of immune tolerance-inducing thymic epithelial cell development by Spi-B-mediated negative feedback regulation. The Journal of experimental medicine 211: 2425–2438

Akiyama T, Shimo Y, Yanai H, Qin J, Ohshima D, Maruyama Y, Asaumi Y, Kitazawa J, Takayanagi H, Penninger JM, Matsumoto M, Nitta T, Takahama Y, Inoue J (2008) The tumor necrosis factor family receptors RANK and CD40 cooperatively establish the thymic medullary microenvironment and self-tolerance. Immunity 29: 423–437

Anderson MS, Venanzi ES, Klein L, Chen Z, Berzins SP, Turley SJ, von Boehmer H, Bronson R, Dierich A, Benoist C, Mathis D (2002) Projection of an immunological self shadow within the thymus by the aire protein. Science 298: 1395–1401

Asano T, Okamoto K, Nakai Y, Tsutsumi M, Muro R, Suematsu A, Hashimoto K, Okamura T, Ehata S, Nitta T, Takayanagi H (2019) Soluble RANKL is physiologically dispensable but accelerates tumour metastasis to bone. Nat Metab 1: 868–875

Baba T, Nakamoto Y, Mukaida N (2009) Crucial contribution of thymic Sirp alpha+ conventional dendritic cells to central tolerance against blood-borne antigens in a CCR2-dependent manner. Journal of immunology 183: 3053–3063

Baran-Gale J, Morgan MD, Maio S, Dhalla F, Calvo-Asensio I, Deadman ME, Handel AE, Maynard A, Chen S, Green F, Sit RV, Neff NF, Darmanis S, Tan W, May AP, Marioni JC, Ponting CP, Holl√§nder GA (2020) Ageing compromises mouse thymus function and remodels epithelial cell differentiation. Elife 9

Barnden MJ, Allison J, Heath WR, Carbone FR (1998) Defective TCR expression in transgenic mice constructed using cDNA-based alpha- and beta-chain genes under the control of heterologous regulatory elements. Immunol Cell Biol 76: 34–40

Bornstein C, Nevo S, Giladi A, Kadouri N, Pouzolles M, Gerbe F, David E, Machado A, Chuprin A, Toth B, Goldberg O, Itzkovitz S, Taylor N, Jay P, Zimmermann VS, Abramson J, Amit I (2018) Single-cell mapping of the thymic stroma identifies IL-25-producing tuft epithelial cells. Nature 559: 622–626

Burkly L, Hession C, Ogata L, Reilly C, Marconi LA, Olson D, Tizard R, Cate R, Lo D (1995) Expression of relB is required for the development of thymic medulla and dendritic cells. Nature 373: 531–536

Chuprin A, Avin A, Goldfarb Y, Herzig Y, Levi B, Jacob A, Sela A, Katz S, Grossman M, Guyon C, Rathaus M, Cohen HY, Sagi I, Giraud M, McBurney MW, Husebye ES, Abramson J (2015) The deacetylase Sirt1 is an essential regulator of Aire-mediated induction of central immunological tolerance. Nature immunology 16: 737–745

Cowan JE, Baik S, McCarthy NI, Parnell SM, White AJ, Jenkinson WE, Anderson G (2018) Aire controls the recirculation of murine Foxp3(+) regulatory T-cells back to the thymus. European journal of immunology 48: 844–854

Derbinski J, Gabler J, Brors B, Tierling S, Jonnakuty S, Hergenhahn M, Peltonen L, Walter J, Kyewski B (2005) Promiscuous gene expression in thymic epithelial cells is regulated at multiple levels. J Exp Med 202: 33–45

Derbinski J, Schulte A, Kyewski B, Klein L (2001) Promiscuous gene expression in medullary thymic epithelial cells mirrors the peripheral self. Nat Immunol 2: 1032–1039

Dhalla F, Baran-Gale J, Maio S, Chappell L, Hollander GA, Ponting CP (2020) Biologically indeterminate yet ordered promiscuous gene expression in single medullary thymic epithelial cells. The EMBO journal 39: e101828

Gabler J, Arnold J, Kyewski B (2007) Promiscuous gene expression and the developmental dynamics of medullary thymic epithelial cells. Eur J Immunol 37: 3363–3372

Giraud M, Yoshida H, Abramson J, Rahl PB, Young RA, Mathis D, Benoist C (2012) Aire unleashes stalled RNA polymerase to induce ectopic gene expression in thymic epithelial cells. Proceedings of the National Academy of Sciences of the United States of America 109: 535–540

Goldfarb Y, Kadouri N, Levi B, Sela A, Herzig Y, Cohen RN, Hollenberg AN, Abramson J (2016) HDAC3 Is a Master Regulator of mTEC Development. Cell Rep 15: 651–665

Gray D, Abramson J, Benoist C, Mathis D (2007) Proliferative arrest and rapid turnover of thymic epithelial cells expressing Aire. J Exp Med 204: 2521–2528

Gray DH, Seach N, Ueno T, Milton MK, Liston A, Lew AM, Goodnow CC, Boyd RL (2006) Developmental kinetics, turnover, and stimulatory capacity of thymic epithelial cells. Blood 108: 3777–3785

Haljasorg U, Dooley J, Laan M, Kisand K, Bichele R, Liston A, Peterson P (2017) Irf4 Expression in Thymic Epithelium Is Critical for Thymic Regulatory T Cell Homeostasis. Journal of immunology 198: 1952–1960

Handel AE, Shikama-Dorn N, Zhanybekova S, Maio S, Graedel AN, Zuklys S, Ponting CP, Hollander GA (2018) Comprehensively Profiling the Chromatin Architecture of Tissue Restricted Antigen Expression in Thymic Epithelial Cells Over Development. Front Immunol 9: 2120

Hikosaka Y, Nitta T, Ohigashi I, Yano K, Ishimaru N, Hayashi Y, Matsumoto M, Matsuo K, Penninger JM, Takayanagi H, Yokota Y, Yamada H, Yoshikai Y, Inoue J, Akiyama T, Takahama Y (2008) The cytokine RANKL produced by positively selected thymocytes fosters medullary thymic epithelial cells that express autoimmune regulator. Immunity 29: 438–450

Hu Z, Lancaster JN, Sasiponganan C, Ehrlich LI (2015) CCR4 promotes medullary entry and thymocyte-dendritic cell interactions required for central tolerance. The Journal of experimental medicine 212: 1947–1965

Irla M, Guerri L, Guenot J, Serge A, Lantz O, Liston A, Imhof BA, Palmer E, Reith W (2012) Antigen recognition by autoreactive cd4(+) thymocytes drives homeostasis of the thymic medulla. PLoS One 7: e52591

Irla M, Hollander G, Reith W (2010) Control of central self-tolerance induction by autoreactive CD4+ thymocytes. Trends in immunology 31: 71–79

Irla M, Hugues S, Gill J, Nitta T, Hikosaka Y, Williams IR, Hubert FX, Scott HS, Takahama Y, Hollander GA, Reith W (2008) Autoantigen-specific interactions with CD4+ thymocytes control mature medullary thymic epithelial cell cellularity. Immunity 29: 451–463

Kadouri N, Nevo S, Goldfarb Y, Abramson J (2019) Thymic epithelial cell heterogeneity: TEC by TEC. Nature reviews Immunology

Klein L, Kyewski B, Allen PM, Hogquist KA (2014) Positive and negative selection of the T cell repertoire: what thymocytes see (and don’t see). Nature reviews Immunology 14: 377–391

Klein L, Robey EA, Hsieh CS (2019) Central CD4(+) T cell tolerance: deletion versus regulatory T cell differentiation. Nature reviews Immunology 19: 7–18

Koh AS, Miller EL, Buenrostro JD, Moskowitz DM, Wang J, Greenleaf WJ, Chang HY, Crabtree GR (2018) Rapid chromatin repression by Aire provides precise control of immune tolerance. Nat Immunol 19: 162–172

Kurts C, Heath WR, Carbone FR, Allison J, Miller JF, Kosaka H (1996) Constitutive class I-restricted exogenous presentation of self antigens in vivo. J Exp Med 184: 923–930

Kyewski B, Klein L (2006) A central role for central tolerance. Annu Rev Immunol 24: 571–606

LeibundGut-Landmann S, Waldburger JM, Reis e Sousa C, Acha-Orbea H, Reith W (2004) MHC class II expression is differentially regulated in plasmacytoid and conventional dendritic cells. Nat Immunol 5: 899–908

Lkhagvasuren E, Sakata M, Ohigashi I, Takahama Y (2013) Lymphotoxin beta receptor regulates the development of CCL21-expressing subset of postnatal medullary thymic epithelial cells. Journal of immunology 190: 5110–5117

Lomada D, Liu B, Coghlan L, Hu Y, Richie ER (2007) Thymus medulla formation and central tolerance are restored in IKKalpha^-/-^mice that express an IKKalpha transgene in keratin 5+ thymic epithelial cells. J Immunol 178: 829–837

Lopes N, Charaix J, Cedile O, Serge A, Irla M (2018) Lymphotoxin alpha fine-tunes T cell clonal deletion by regulating thymic entry of antigen-presenting cells. Nat Commun 9: 1262

Lopes N, Serge A, Ferrier P, Irla M (2015) Thymic Crosstalk Coordinates Medulla Organization and T-Cell Tolerance Induction. Front Immunol 6: 365

Lopes N, Vachon H, Marie J, Irla M (2017) Administration of RANKL boosts thymic regeneration upon bone marrow transplantation. EMBO Mol Med

Madsen L, Labrecque N, Engberg J, Dierich A, Svejgaard A, Benoist C, Mathis D, Fugger L (1999) Mice lacking all conventional MHC class II genes. Proceedings of the National Academy of Sciences of the United States of America 96: 10338–10343

Metzger TC, Khan IS, Gardner JM, Mouchess ML, Johannes KP, Krawisz AK, Skrzypczynska KM, Anderson MS (2013) Lineage tracing and cell ablation identify a post-Aire-expressing thymic epithelial cell population. Cell Rep 5: 166–179

Michel C, Miller CN, Kuchler R, Brors B, Anderson MS, Kyewski B, Pinto S (2017) Revisiting the Road Map of Medullary Thymic Epithelial Cell Differentiation. Journal of immunology 199: 3488–3503

Miller CN, Proekt I, von Moltke J, Wells KL, Rajpurkar AR, Wang H, Rattay K, Khan IS, Metzger TC, Pollack JL, Fries AC, Lwin WW, Wigton EJ, Parent AV, Kyewski B, Erle DJ, Hogquist KA, Steinmetz LM, Locksley RM, Anderson MS (2018) Thymic tuft cells promote an IL-4-enriched medulla and shape thymocyte development. Nature 559: 627–631

Nishikawa Y, Hirota F, Yano M, Kitajima H, Miyazaki J, Kawamoto H, Mouri Y, Matsumoto M (2010) Biphasic Aire expression in early embryos and in medullary thymic epithelial cells before end-stage terminal differentiation. The Journal of experimental medicine 207: 963–971

Org T, Rebane A, Kisand K, Laan M, Haljasorg U, Andreson R, Peterson P (2009) AIRE activated tissue specific genes have histone modifications associated with inactive chromatin. Hum Mol Genet 18: 4699–4710

Otero DC, Baker DP, David M (2013) IRF7-Dependent IFN-beta Production in Response to RANKL Promotes Medullary Thymic Epithelial Cell Development. Journal of immunology

Pezzi N, Assis AF, Cotrim-Sousa LC, Lopes GS, Mosella MS, Lima DS, Bombonato-Prado KF, Passos GA (2016) Aire knockdown in medullary thymic epithelial cells affects Aire protein, deregulates cell adhesion genes and decreases thymocyte interaction. Mol Immunol 77: 157–173

Riemann M, Andreas N, Fedoseeva M, Meier E, Weih D, Freytag H, Schmidt-Ullrich R, Klein U, Wang ZQ, Weih F (2017) Central immune tolerance depends on crosstalk between the classical and alternative NF-kappaB pathways in medullary thymic epithelial cells. J Autoimmun 81: 56–67

Rodrigues PM, Ribeiro AR, Perrod C, Landry JJM, Araujo L, Pereira-Castro I, Benes V, Moreira A, Xavier-Ferreira H, Meireles C, Alves NL (2017) Thymic epithelial cells require p53 to support their long-term function in thymopoiesis in mice. Blood 130: 478–488

Sansom SN, Shikama-Dorn N, Zhanybekova S, Nusspaumer G, Macaulay IC, Deadman ME, Heger A, Ponting CP, Hollander GA (2014) Population and single-cell genomics reveal the Aire dependency, relief from Polycomb silencing, and distribution of self-antigen expression in thymic epithelia. Genome Res

Serge A, Bailly AL, Aurrand-Lions M, Imhof BA, Irla M (2015) For3D: Full Organ Reconstruction in 3D, an Automatized Tool for Deciphering the Complexity of Lymphoid Organs. Journal of immunological methods

Shen H, Ji Y, Xiong Y, Kim H, Zhong X, Jin MG, Shah YM, Omary MB, Liu Y, Qi L, Rui L (2019) Medullary thymic epithelial NF-kB-inducing kinase (NIK)/IKKalpha pathway shapes autoimmunity and liver and lung homeostasis in mice. Proceedings of the National Academy of Sciences of the United States of America 116: 19090–19097

Shinkai Y, Rathbun G, Lam KP, Oltz EM, Stewart V, Mendelsohn M, Charron J, Datta M, Young F, Stall AM, et al. (1992) RAG-2-deficient mice lack mature lymphocytes owing to inability to initiate V(D)J rearrangement. Cell 68: 855–867

Takaba H, Morishita Y, Tomofuji Y, Danks L, Nitta T, Komatsu N, Kodama T, Takayanagi H (2015) Fezf2 Orchestrates a Thymic Program of Self-Antigen Expression for Immune Tolerance. Cell 163: 975–987

Tykocinski LO, Sinemus A, Rezavandy E, Weiland Y, Baddeley D, Cremer C, Sonntag S, Willecke K, Derbinski J, Kyewski B (2010) Epigenetic regulation of promiscuous gene expression in thymic medullary epithelial cells. Proceedings of the National Academy of Sciences of the United States of America 107: 19426–19431

Ucar O, Rattay K (2015) Promiscuous Gene Expression in the Thymus: A Matter of Epigenetics, miRNA, and More? Front Immunol 6: 93

Ueno T, Saito F, Gray DH, Kuse S, Hieshima K, Nakano H, Kakiuchi T, Lipp M, Boyd RL, Takahama Y (2004) CCR7 signals are essential for cortex-medulla migration of developing thymocytes. J Exp Med 200: 493–505

van Ewijk W, Shores EW, Singer A (1994) Crosstalk in the mouse thymus. Immunol Today 15: 214–217

Waldburger JM, Rossi S, Hollander GA, Rodewald HR, Reith W, Acha-Orbea H (2003) Promoter IV of the class II transactivator gene is essential for positive selection of CD4+ T cells. Blood 101: 3550–3559

White AJ, Jenkinson WE, Cowan JE, Parnell SM, Bacon A, Jones ND, Jenkinson EJ, Anderson G (2014) An essential role for medullary thymic epithelial cells during the intrathymic development of invariant NKT cells. Journal of immunology 192: 2659–2666

Wong K, Lister NL, Barsanti M, Lim JM, Hammett MV, Khong DM, Siatskas C, Gray DH, Boyd RL, Chidgey AP (2014) Multilineage potential and self-renewal define an epithelial progenitor cell population in the adult thymus. Cell Rep 8: 1198–1209

Youm YH, Horvath TL, Mangelsdorf DJ, Kliewer SA, Dixit VD (2016) Prolongevity hormone FGF21 protects against immune senescence by delaying age-related thymic involution. Proceedings of the National Academy of Sciences of the United States of America 113: 1026–1031

Zhang B, Wang Z, Ding J, Peterson P, Gunning WT, Ding HF (2006) NF-kappaB2 is required for the control of autoimmunity by regulating the development of medullary thymic epithelial cells. J Biol Chem 281: 38617–38624

Zhu M, Chin RK, Christiansen PA, Lo JC, Liu X, Ware C, Siebenlist U, Fu YX (2006) NF-kappaB2 is required for the establishment of central tolerance through an Aire-dependent pathway. J Clin Invest 116: 2964–2971

Zuklys S, Handel A, Zhanybekova S, Govani F, Keller M, Maio S, Mayer CE, Teh HY, Hafen K, Gallone G, Barthlott T, Ponting CP, Hollander GA (2016) Foxn1 regulates key target genes essential for T cell development in postnatal thymic epithelial cells. Nature immunology 17: 1206–1215

