## Supplemental Figures for "Thymocytes trigger self-antigen-controlling pathways in immature medullary thymic epithelial stages"

**Figure S1.** Gating strategy used to purify mTEC$^{lo}$ cells ……………………….….2

**Figure S2.** Normal total and phosphorylated p65, RelB and Erk1/2 proteins in mTEC$^{lo}$ from ΔCD4 mice ………………………………………………3

**Figure S3.** Impaired TRA expression and cellularity of Fezf2$^{+}$ and DCLK1$^{+}$ mTECs in MHCII$^{-/-}$ mice……………………….…………..….…………………4

**Figure S4.** Normal proliferation of Aire$^{-}$Fezf2$^{+}$ and Aire$^{+}$Fezf2$^{+}$ mTECs in ΔCD4 and mTEC$^{ΔMHCII}$ mice……………………………………………….….…5

**Figure S5**. Reduced post-Aire mTECs in ΔCD4 and mTEC$^{ΔMHCII}$ mice.......……..6

**Figure S6.** Similar levels of total and phosphorylated p65, Erk1/2, p38 and IKKα proteins in mTEC$^{lo}$ from RipmOVAxOTII-*Rag2$^{-/-}$* and OTII-*Rag2$^{-/-}$* mice……………………………..…………………………………………….…7

**Figure S7.** The proliferation of Aire$^{-}$Fezf2$^{+}$ and Aire$^{+}$Fezf2$^{+}$ mTECs is similar in RipmOVAxOTII-*Rag2$^{-/-}$* and OTII-*Rag2$^{-/-}$* mice……………………………….....8

**Figure S8**. Post-Aire mTECs are increased in RipmOVAxOTII-*Rag2$^{-/-}$* compared to OTII-*Rag2$^{-/-}$* mice…………………………………………………………..…9

### Figure S1

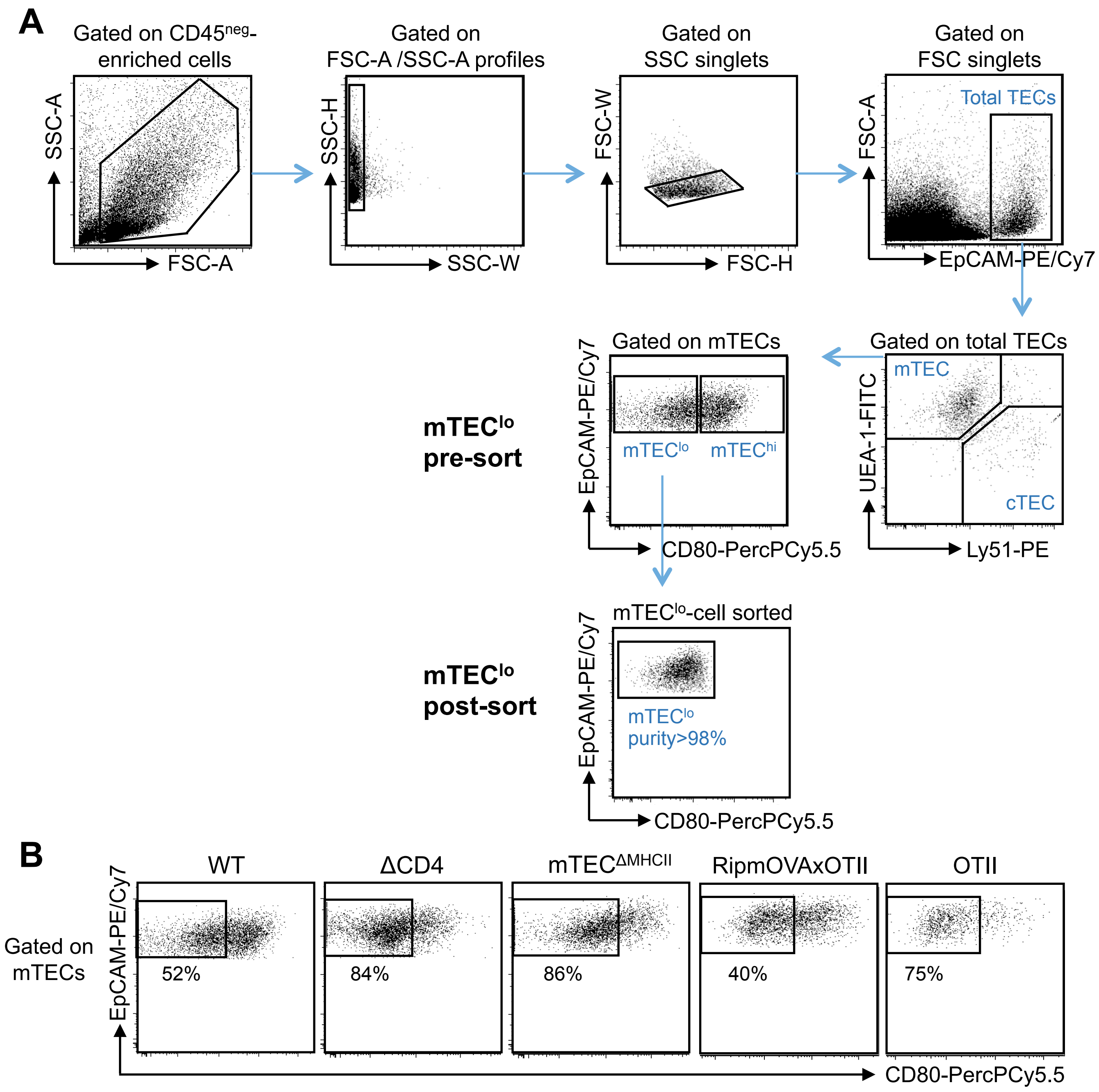


**Figure S1. Gating strategy used to purify mTEC$^{lo}$ cells.**

**(A)** Total TECs were defined as EpCAM$^{+}$ in CD45$^{neg}$-enriched thymic cells by autoMACS and were further divided into mTECs (UEA-1$^{+}$Ly51$^{lo}$) and cTECs (UEA-1$^{-}$Ly51$^{hi}$). mTEC$^{lo}$ were identified and sorted based on low/intermediate level of the CD80 co-stimulatory molecule. The purity of sorted mTEC$^{lo}$ was >98%.
**(B)** Gates used to sort mTEC$^{lo}$ from WT, ΔCD4, mTEC$^{ΔMHCII}$, RipmOVAxOTII-*Rag2*$^{-/-}$ and OTII-*Rag2*$^{-/-}$ mice. This gating strategy was used to sort or analyse mTEC$^{lo}$ in Fig. 1; 2B-K; 3B,C; 4; 5F,G; 6 and in Fig S2; S3; S4; S6 and S7.

### Figure S2

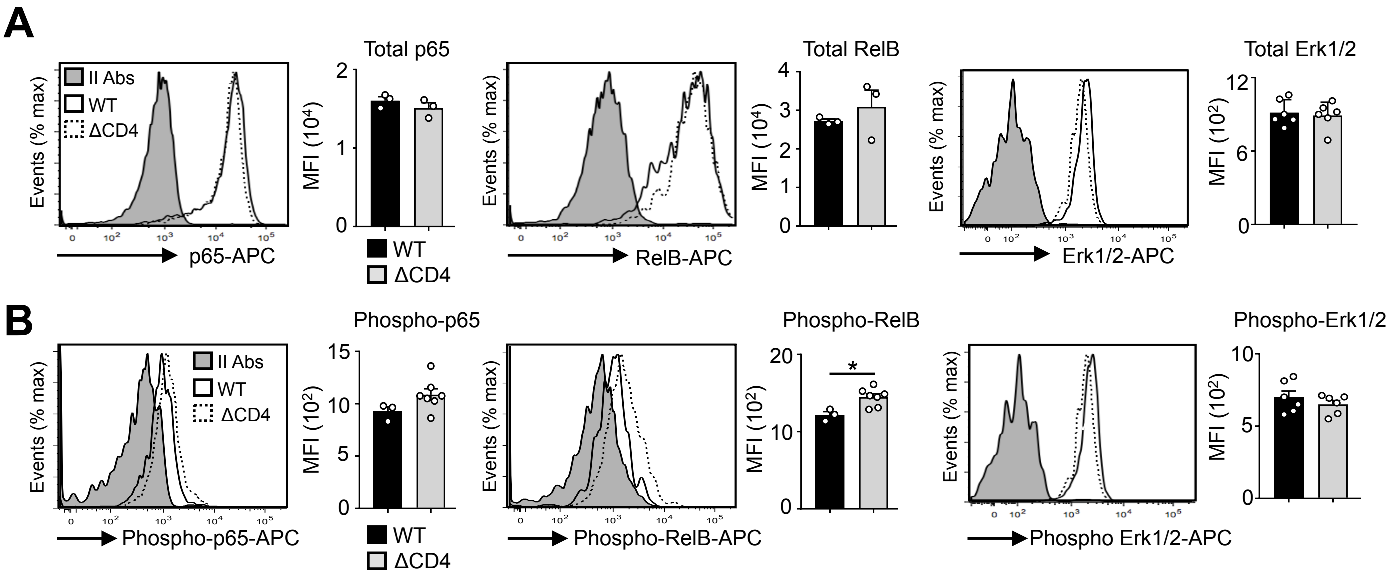


**Figure S2. Normal total and phosphorylated p65, RelB and Erk1/2 proteins in mTEC$^{lo}$ from ΔCD4 mice.**

Total p65, RelB and Erk1/2 **(A)** and phospho-p65 (Ser536), phospho-RelB (Ser552) and phospho-Erk1/2 (Thr202/Tyr204) **(B)** proteins were analyzed by flow cytometry in mTEC$^{lo}$ of WT and ΔCD4 mice. Histograms show the MFI. II Abs: Secondary antibodies. Data are representative of 2 independent experiments (n=2-3 mice per group and experiment). Error bars show mean±SEM, *p<0.05 using the Mann-Whitney test.

### Figure S3

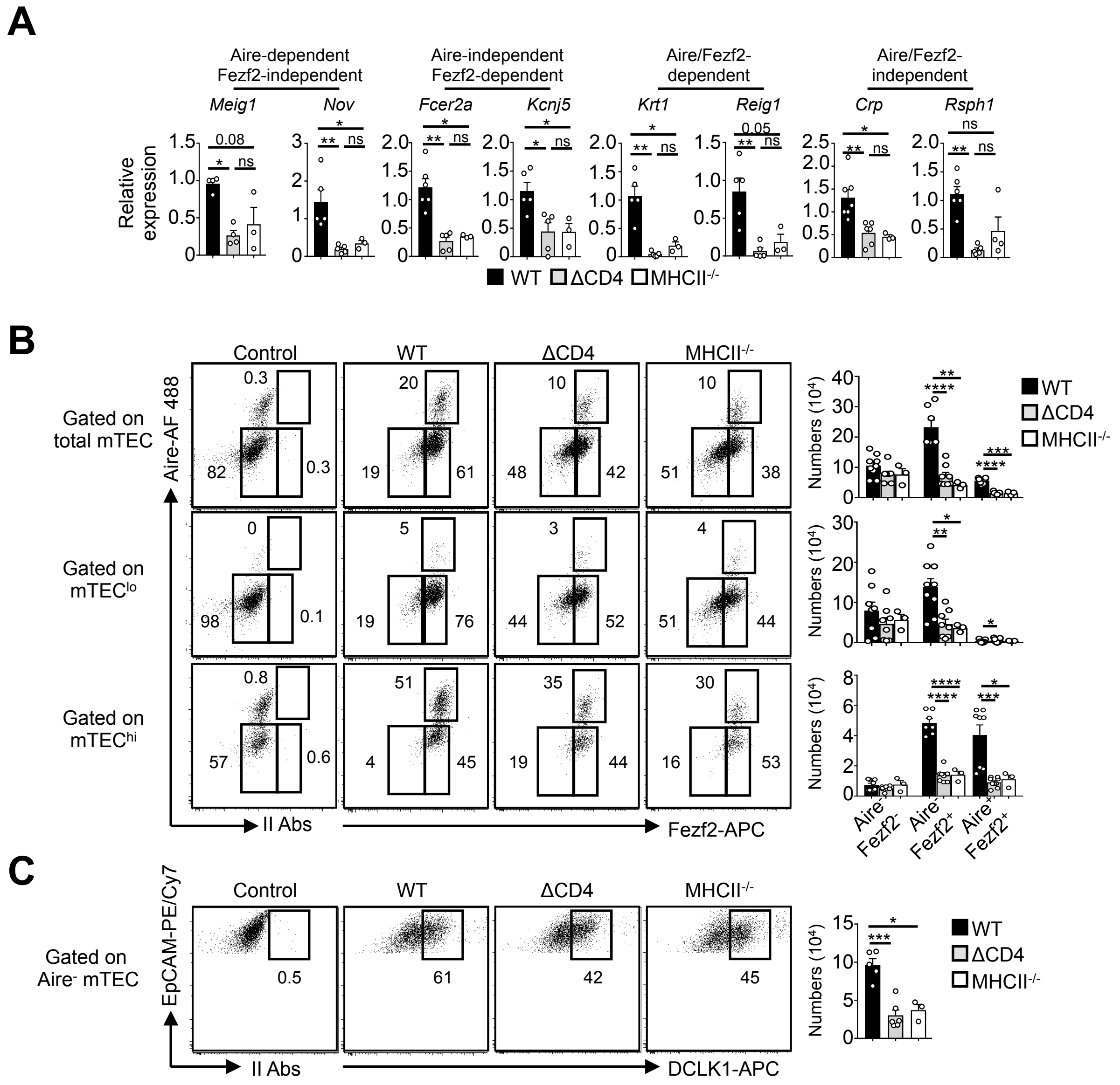


**Figure S3. Impaired TRA expression and cellularity of Fezf2$^+$ and DCLK1$^+$ mTECs in MHCII$^{-/-}$ mice.**

**(A)** The expression of Aire-dependent (*Meig1, Nov*), Fezf2-dependent (*Fcer2a, Kcnj5*), Aire/Fezf2-dependent (*Krt1, Reig1*) and Aire/Fezf2-independent (*Crp, Rsph1*) TRAs was measured by qPCR in purified mTEC$^{lo}$ from WT (n=3), ΔCD4 (n=3) and MHCII$^{-/-}$ (n=3) mice.
**(B)** Flow cytometry profiles and numbers of Aire$^-$Fezf2$^-$, Aire$^-$Fezf2$^+$ and Aire$^+$Fezf2$^+$ cells in total mTECs, mTEC$^{lo}$ and mTEC$^{hi}$ from WT, ΔCD4 and MHCII$^{-/-}$ mice. II Abs: secondary antibodies. Data are representative of 2 independent experiments (n=2-4 mice per group).
**(C)** Flow cytometry profiles and numbers of DCKL1$^+$ cells in Aire$^-$ mTECs. Data are representative of 2 independent experiments (n = 2-3 mice per group). Error bars show mean ± SEM, $*p < 0.05$, $**p<0.01$, $***p<0.001$, $****p<0.0001$ using the Mann-Whitney test.

### Figure S4

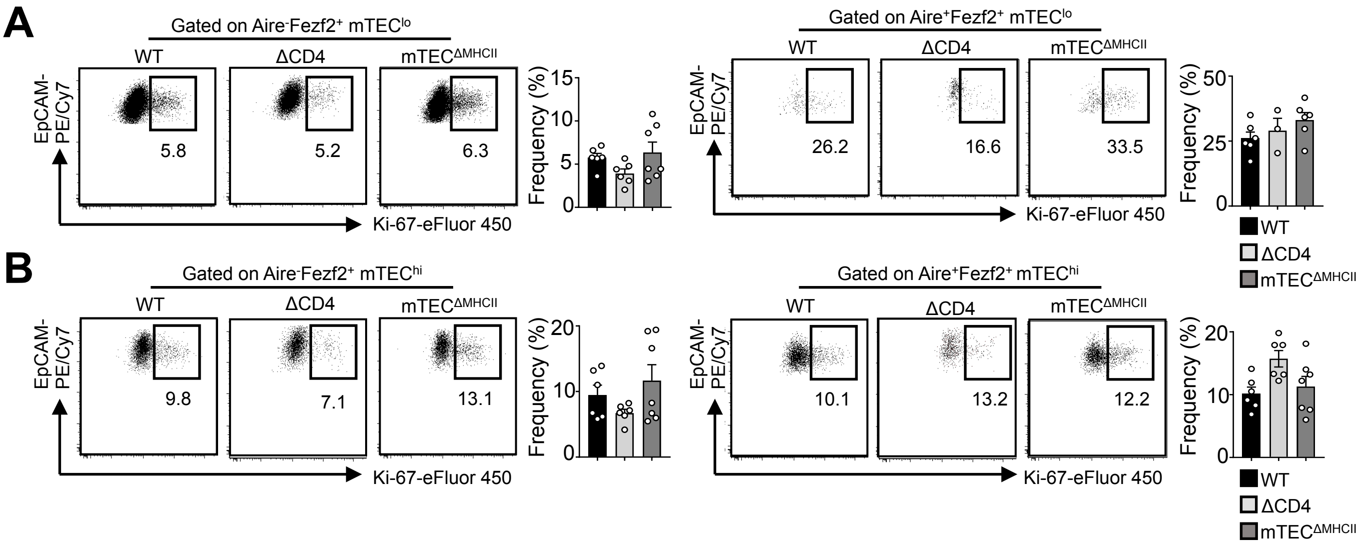


**Figure S4. Normal proliferation of Aire⁻Fezf2⁺ and Aire⁺Fezf2⁺ mTECs in ΔCD4 and mTEC$^{\Delta MHCII}$ mice.**

**(A,B)** Flow cytometry profiles and frequencies of proliferating Ki-67$^{+}$ Aire$^{-}$Fezf2$^{+}$ and Aire$^{+}$Fezf2$^{+}$ cells in mTEC$^{lo}$ **(A)** and mTEC$^{hi}$ **(B)** from WT, ΔCD4 and mTEC$^{\Delta MHCII}$ mice. Data are representative of 2 independent experiments (n=3-4 mice per group and experiment). Error bars show mean±SEM.

### Figure S5

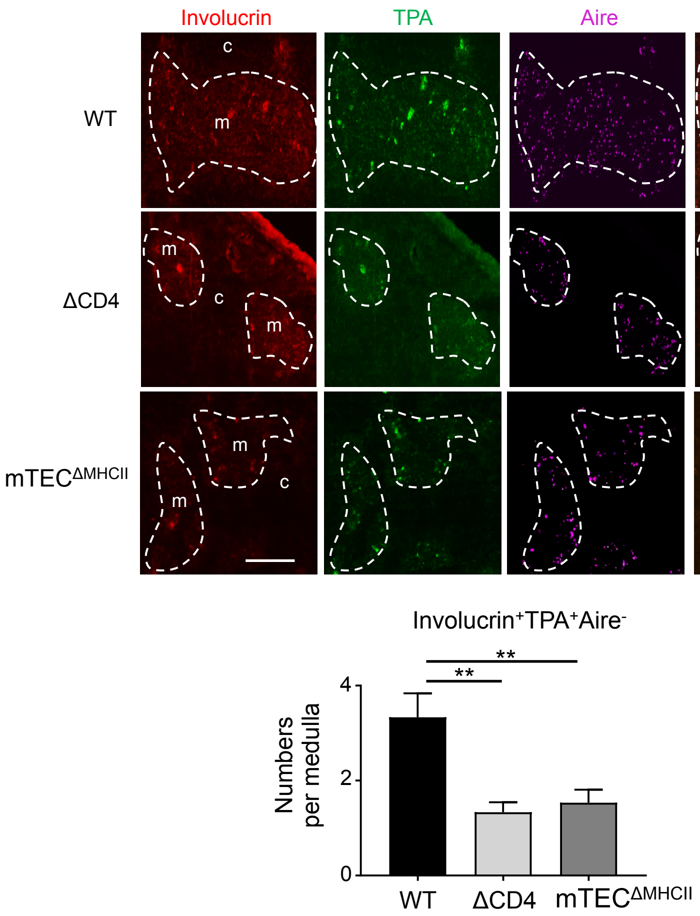


**Figure S5. Reduced post-Aire mTECs in ΔCD4 and mTEC$^{\Delta MHCII}$ mice.**

Representative thymic sections stained with antibodies against involucrin (red), TPA (green) and Aire (magenta). c and m denotes the cortex and the medulla, respectively. The graph shows the number of Involucrin$^{+}$TPA$^{+}$Aire$^{-}$ cells per medulla. 15 medullas derived from 2 WT, 2 ΔCD4 and 2 mTEC$^{\Delta MHCII}$ mice were quantified, respectively. Scale bar: 200 μm. Error bars show mean±SEM, **p<0.01 using unpaired Student's t-test.

### Figure S6

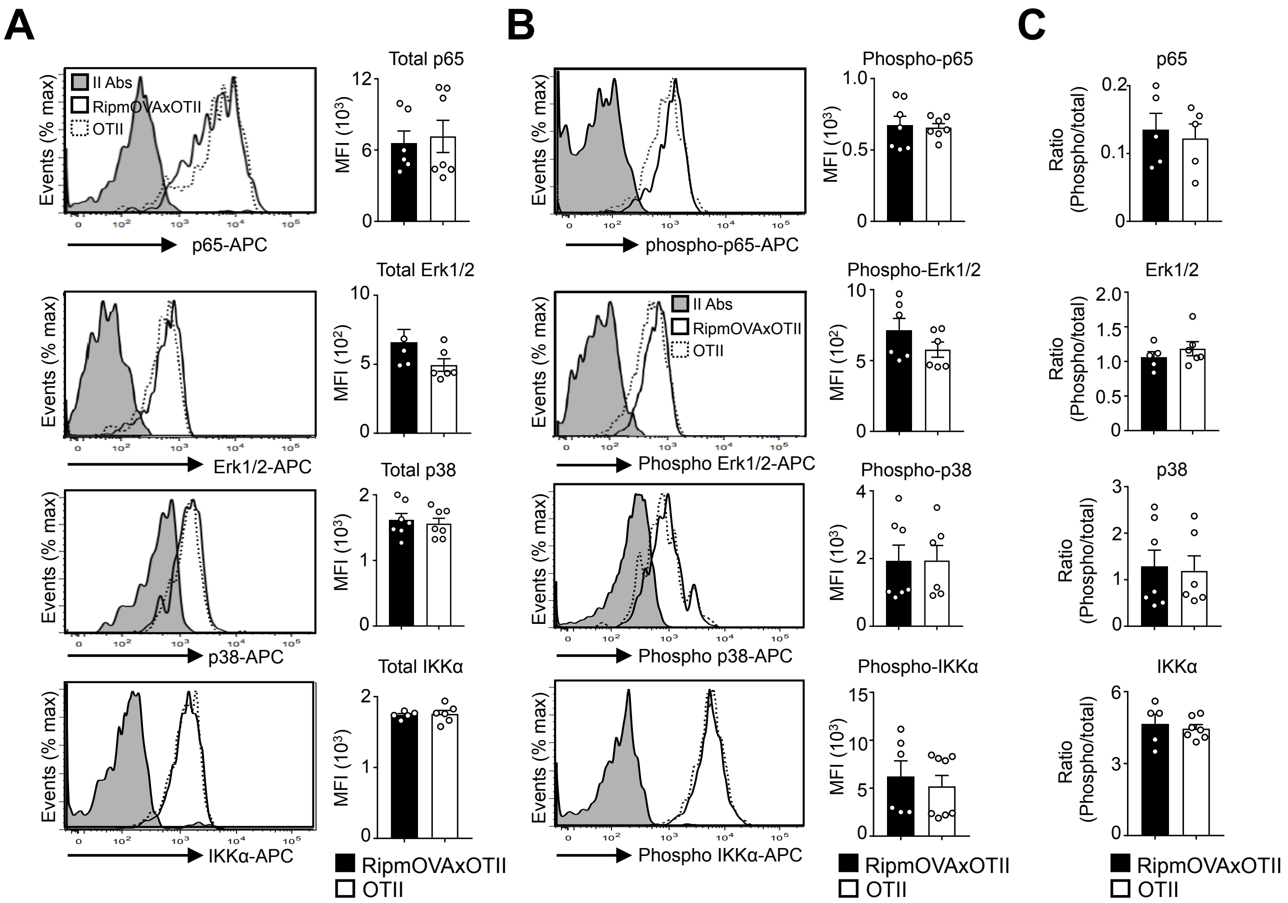


**Figure S6. Similar levels of total and phosphorylated p65, Erk1/2, p38 and IKKα proteins in mTEC$^{lo}$ from RipmOVAxOTII-*Rag2*$^{-/-}$ and OTII-*Rag2*$^{-/-}$ mice.**

**(A,B)** Total p65, Erk1/2, p38 and IKKα **(A)** and phospho-p65 (Ser536), phospho-Erk1/2 MAPK (Thr202/Tyr204), phospho-p38 MAPK (Thr180/Tyr182) and phospho-IKKα(Ser180)/ IKKβ(Ser181) **(B)** proteins were analyzed by flow cytometry in mTEC$^{lo}$ of RipmOVAxOTII-*Rag2*$^{-/-}$ and OTII-*Rag2*$^{-/-}$ mice. Histograms show the MFI.
**(C)** Histograms represent the ratio of phospho-p65 (Ser536) to p65, phospho-Erk1/2 MAPK (Thr202/Tyr204) to Erk1/2, phospho-p38 MAPK (Thr180/Tyr182) to p38 and phospho-IKKα (Ser180)/IKKβ(Ser181) to IKKα. II Abs: secondary antibodies. Data are representative of 2 independent experiments (n=2-4 mice per group and experiment). Error bars show mean±SEM.

### Figure S7

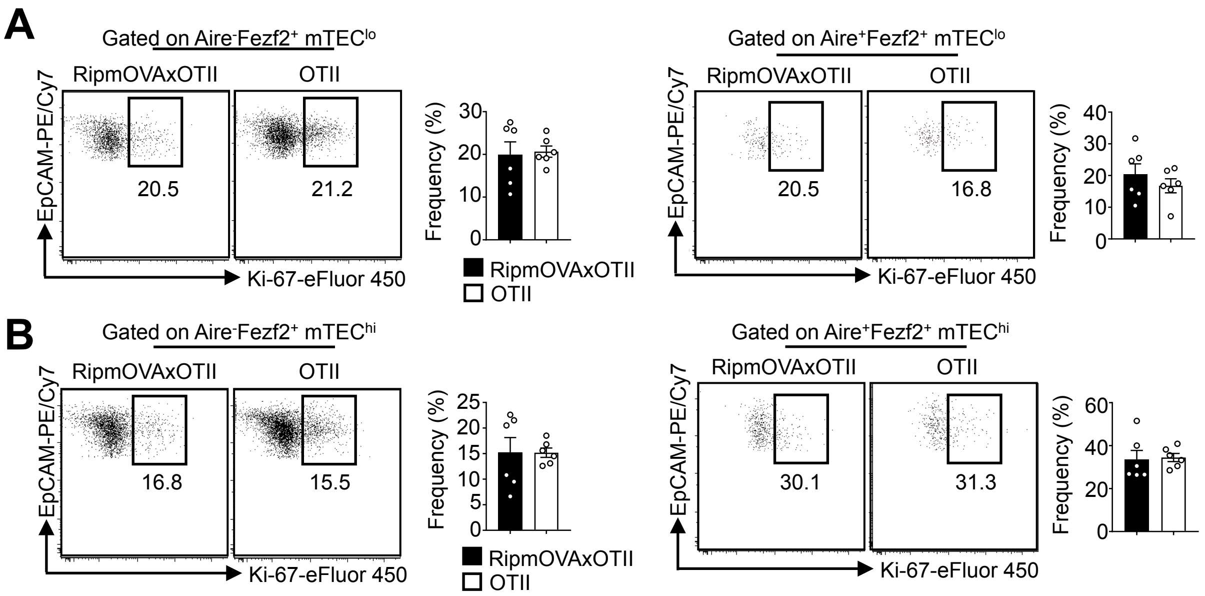


**Figure S7. The proliferation of Aire$^-$Fezf2$^+$ and Aire$^+$Fezf2$^+$ mTECs is similar in RipmOVAxOTII-*Rag2$^{-/-}$* and OTII-*Rag2$^{-/-}$* mice.**

**(A,B)** Flow cytometry profiles and frequencies of proliferating Ki-67$^+$ Aire$^-$Fezf2$^+$ and Aire$^+$Fezf2$^+$ in mTEC$^{lo}$ **(A)** and mTEC$^{hi}$ **(B)** from RipmOVAxOTII-*Rag2$^{-/-}$* and OTII-*Rag2$^{-/-}$* mice. Data are representative of 2 independent experiments (n=3 mice per group and experiment). Error bars show mean±SEM.

### Figure S8

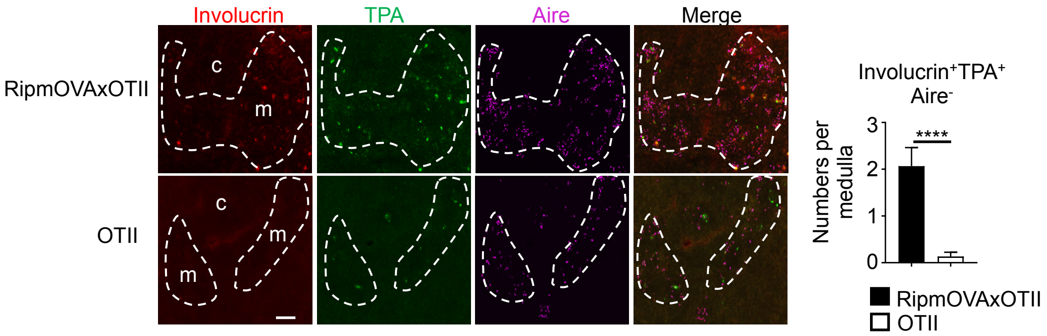


**Figure S8. Post-Aire mTECs are increased in RipmOVAxOTII-*Rag2*$^{-/-}$ compared to OTII-*Rag2*$^{-/-}$ mice.**

Representative thymic sections from RipmOVAxOTII-*Rag2*$^{-/-}$ and OTII-*Rag2*$^{-/-}$ mice stained with antibodies against involucrin (red), TPA (green) and Aire (magenta). c and m denotes the cortex and the medulla, respectively. The graph shows the number of Involucrin$^{+}$TPA$^{+}$Aire$^{-}$ cells per medulla. 15 medullas derived from 2 RipmOVAxOTII-*Rag2*$^{-/-}$ and 2 OTII-*Rag2*$^{-/-}$ mice were quantified, respectively. Scale bar: 200 μm. Error bars show mean ± SEM, ****p<0.0001 using unpaired Student's t-test.
